# Episodic memory adaptively expands retrieval through typicality-guided transformation while concrete details remain accessible

**DOI:** 10.64898/2026.09.24.754119

**Authors:** Mattia Delmarco, Carlos González-García, Juan Linde-Domingo

## Abstract

Memory faces a fundamental challenge. It has to become flexible enough to be accessed by novel cues without losing the specific information that defines each individual episode. Whether memory transformation solves this problem by sacrificing details or by adaptively expanding retrieval while preserving them is unclear. We tested this by manipulating the match between a memory and its retrieval cue. Participants retrieved object associations after 6, 24 and 48 hours using either the original encoded image or novel exemplars that were more typical or more distinctive of the same object category. Retrieval initially benefited from the originally encoded image, but this advantage was reduced across delay. By 48 hours, a never-seen, more typical exemplar triggered retrieval as effectively as the original one, while the same convergence was not evident for less-typical exemplars. Crucially, such cue expansion was not accompanied by a loss of perceptual specificity. Participants retained access to the originally encoded exemplar across delays, while errors increasingly favoured more typical alternatives. At the item level, stronger episodic memories predicted better access to perceptual detail, but did not predict the tendency of errors for category-like information, which instead increased with delay. This dissociation suggests that memory generalisation and the continued accessibility of visual detail reflect partly separable transformations rather than opposite ends of a single continuum.

## Introduction

In The Sopranos, Tony Soprano commissions a painting of himself with the racehorse Pie-O-My. However, after receiving the painting, Tony decides that he wants to get rid of it and asks his associates to destroy it. Instead, his friend Paulie secretly keeps the painting, modifies it, and hangs it in his own home. Crucially, Paulie changes the image: Tony is no longer depicted as in the original portrait, but is transformed into a war-general-like figure. Despite the passage of time and these changes, Tony immediately recognises the altered work when he later sees it at Paulie’s house. This apparently simple recognition requires more than a vague sense of familiarity. It captures a fundamental tension. On one hand, Tony must retain concrete and specific perceptual information about the original image, such as the horse and the composition. On the other hand, he also needs to flexibly generalise across the changes in order to recognise the modified figure as himself.

The scene therefore illustrates a central property of adaptive memory: retrieval must tolerate changes in how it is cued while preserving the perceptual details needed to identify the original episode. How human cognition resolves the trade-off between generalisation and specificity is a central question across the cognitive, neural, and computational sciences (Kumaran et al., 2016; Schacter et al., 2011; Spens & Burgess, 2024; Zeithamova & Bowman, 2020). Human memory offers a particularly tractable scenario to study this question, because the balance between concrete and gist-like content is widely observed to shift over time. In particular, experiences are initially rich in episodic and concrete perceptual details, but over hours and days they appear to become more schematic and more like the categories to which they belong (Bartlett, 1932; Lifanov et al., 2021; Moscovitch et al., 2016; Sekeres et al., 2018; Zeng et al., 2021). What remains unclear is what this gist-like shift actually reflects. Does memory become more schematic because perceptual information is no longer accessible, or does retrieval become increasingly guided by category structure even while perceptual detail remains available if needed (Robin & Moscovitch, 2017; Yonelinas et al., 2019)?

This distinction is fundamental to understanding how adaptive our memory system is. On one hand, if specific content is eroded, then there is a cost of generalisation: the system trades fidelity for flexibility. On the other hand, if specific content is preserved but progressively transformed to better match the learned category structure, the picture is fundamentally different: the system retains its capacity for fine-grained access when it is needed, while gaining the potential advantages of generalisation. This would allow us to make adaptive use of novel cues to retrieve our memories without losing the discrimination of concrete details. These two possibilities make different predictions about the architecture of adaptive cognition, and converge with parallel debates in machine learning about how artificial systems can consolidate knowledge without catastrophically overwriting it (Van De Ven et al., 2020).

Different influential theoretical perspectives favour the more adaptive possibility. Trace transformation theory and complementary learning systems propose that detailed and gist-like representations co-exist, with their relative dominance shifting over consolidation through reweighting rather than replacement (McClelland et al., 1995; Moscovitch et al., 2016; Sekeres et al., 2018). Consistent with this idea, neural representations can simultaneously support integration across related experiences while preserving distinctions between individual events, and semantic representations can transform over time while perceptual representations remain comparatively preserved (Audrain & McAndrews, 2022; Krenz et al., 2023; Schlichting et al., 2015; Tompary & Davachi, 2017). Bayesian and schema-based accounts of reconstructive memory complementarily predict that as episodic precision is reduced, retrieval is increasingly weighted by category-level priors, producing systematic biases towards typical exemplars (Hemmer & Steyvers, 2009a; Huttenlocher et al., 2000; Kerrén et al., 2024; Persaud & Hemmer, 2016; Ramey, 2026; Tandoc et al., 2024; Tompary & Thompson-Schill, 2021). Together, these frameworks make a strong joint prediction that has not yet been directly tested. If consolidation reweights rather than replaces content, then the transformed trace should display three behavioural signatures simultaneously: (i) it should *broaden the set of cues that can effectively trigger retrieval*, allowing never-encoded but more category-typical exemplars to function as well as the originally encoded cue; (ii) *fine-grained perceptual information should remain behaviourally accessible*; and (iii) when retrieval of concrete details fails, the resulting reconstruction should be *systematically pulled towards category-typical content*, reflecting the increased contribution of category-level structure. The first of these signatures (input-side generalisation) is the central claim of the Dynamic-Cueing hypothesis (Linde-Domingo & Kerrén, 2025), which formalises the idea that retrieval success depends on cue alignment with the current representational state of the trace rather than with the original encoding episode (cf. Tulving & Thomson, 1973). Together, input-side generalisation, output-side preservation, and reconstruction-side typicality drift would constitute three observable features of an adaptive transformation. However, no study has shown them within a single design.

Existing work has approached these predictions separately. Berens, Richards, and Horner (2020) demonstrated that forgetting predominantly involves losses of accessibility rather than precision, but did not test whether errors drift towards category-typical content or whether the set of effective cues broadens. Bayesian-prior studies have demonstrated category-driven distortion, but without manipulating consolidation delay across days (Hemmer & Steyvers, 2009a, 2009b; Kerrén et al., 2024; Huttenlocher et al., 2000). Trace-transformation studies have demonstrated time-dependent neural shifts towards shared, schematic representations, but have not behaviourally separated availability from the constraint regime governing retrieval (Audrain & McAndrews, 2022; Krenz et al., 2023; Tompary & Davachi, 2017). Lifanov et al. (2021), building on the conceptual-before-perceptual reconstruction gradient identified by Linde-Domingo, Treder, Kerrén, and Wimber (2019), showed that the perceptual–conceptual reaction-time gap grows over a 48-hour delay, consistent with semanticisation, but could not adjudicate whether perceptual content was less available or simply less prioritised at retrieval, and was uninformative about both the structure of recall errors and the cues that trigger retrieval. The same ambiguity applies to the reduced use of perceptual features after consolidation (Tarder-Stoll et al., 2024), since less used does not directly reflect unavailability. Relatedly, Heinen et al. (2025) showed that multiple formats coexist within a memory trace, and that targeted memory reactivation during sleep can selectively strengthen whichever format was task-relevant at encoding while leaving the other largely intact. This set of results has also been reflected by a theoretical tension recently articulated by Horner (2026), who argues that the components conflated in systems-consolidation accounts could be mechanistically independent and should be treated as separable dimensions through which a memory can move along non-unidirectional trajectories. On this view, the loss of episodic specificity that often co-occurs with consolidation need not be a consequence of consolidation itself; the two are temporally correlated but causally distinct, and the framework explicitly calls for experimental designs capable of dissociating them. What is missing, then, is a single behavioural paradigm in which input-side cue generalisation, output-side perceptual preservation, and reconstruction-side typicality drift can be measured jointly across a graded delay manipulation, allowing the multiple signatures of trace transformation to be assessed not as facets of a single graded process but as potentially independent trajectories whose joint pattern characterises the architecture of adaptive memory change.

Here, we address these questions in a pre-registered experiment using typicality-graded object stimuli drawn from 112 concepts of the THINGS database (Hebart et al., 2019). Participants learned associations between object exemplars (Fig. 1a) and were later tested across different delays (Fig. 1b) using complementary measures of category-level retrieval, semantic recall, and perceptual discrimination (Fig. 1d). Critically, retrieval cues could be the originally encoded image of a concept (e.g. a chair) or two never-encoded exemplars from the same concept: a more typical chair or a less typical one (Fig. 1c). This design allowed us to test three core predictions, each corresponding to one signature of trace transformation. First, if the transformed trace broadens the set of cues that can effectively trigger retrieval, the privileged status of the originally encoded cue should diminish over time relative to more typical, never-encoded cues. Second, if perceptual content remains available even as retrieval becomes more schema-guided, perceptual discrimination should remain robustly above chance across delays, including 48 hours after encoding. Third, if retrieval is increasingly guided by conceptual structure when the precise trace cannot be accessed, recognition errors should show a systematic and growing bias towards typical concept exemplars. Critically, by measuring all three signatures we could assess whether they constitute facets of a single adaptive transformation or independent processes that scale with delay through separate mechanisms. Such a distinction has implications for understanding whether consolidation operates over a unified trace or over partially separable traces of the same memory.

**Fig. 1.**
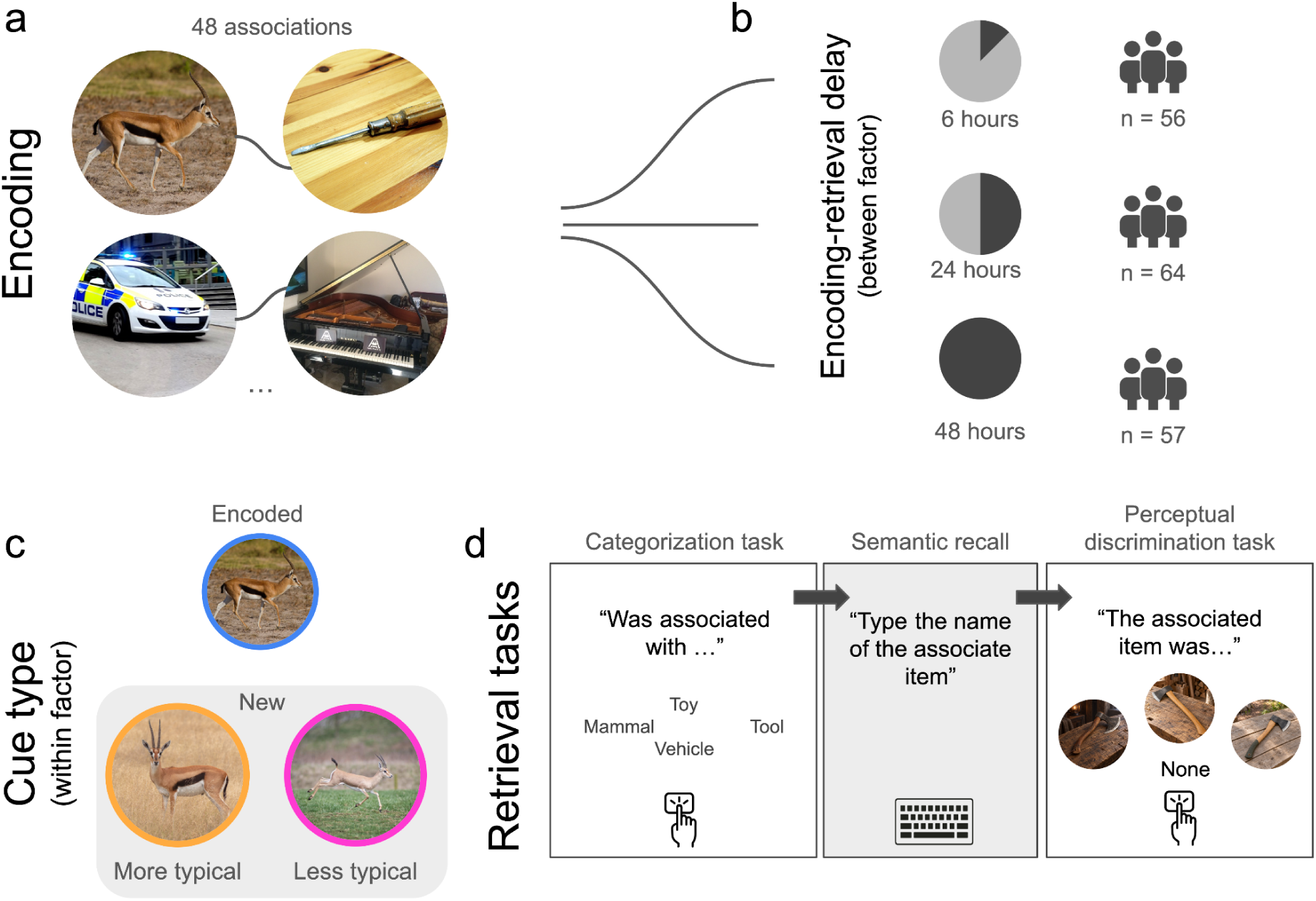
Experimental paradigm. **(a)** Encoding. Participants studied 48 pairs of images and were instructed to form a vivid interactive mental image linking each pair. Each pair remained on screen for a minimum of 2 s and a maximum of 10 s (self-paced), separated by a 500–1000 ms jittered inter-trial interval. The full list was presented across two encoding loops to ensure stable learning. **(b)** Encoding-to-retrieval delay (between-subjects factor). Participants were assigned to one of three retention intervals: 6 h (*n* = 56), 24 h (*n* = 64), or 48 h (*n* = 57). **(c)** Cue typicality (within-subjects factor). For each cue concept, three perceptual variants were drawn from continuous normative typicality ratings obtained in a separate large-scale rank-based judgment study: a medium exemplar (closest to the concept mean), which served as the originally encoded cue at study, and two never-encoded variants: a more typical exemplar and a less typical exemplar. At test, three types of cues were used for each retrieval trial: the originally encoded cue, or two never-encoded variants including a more and a less typical exemplar from the encoded concept. **(d)** Retrieval tasks (fixed order). Each pair was probed using three complementary measures. In *the categorisation task*, participants identified the category of the associated target via one of four labelled response options (e.g., Mammal, Toy, Vehicle, Tool) with key-to-label mapping randomized across trials. In the *semantic recall task*, participants typed the name of the target object, with an auto-suggest tool supporting spelling accuracy. In the perceptual discrimination task, participants selected the associated target from four options comprising the three perceptual variants (more typical, encoded, less typical) of the target plus a “none of these” option. On 25% of trials (Target-absent trials), the target and its variants were replaced by an object from a never-encoded concept, for which “none of these” was the correct response. Each trial began with a jittered fixation (500–1000 ms) and ended with a confidence rating on a 0–100 slider.

We found that all three signatures emerged. The privileged status of the originally encoded cue weakened over time, with never-encoded but more typical cues becoming as effective as the encoded ones at long delays. Perceptual discrimination remained robustly above chance throughout, indicating that fine-grained content remained accessible across delays. Finally, errors in the perceptual discrimination task became increasingly biased towards category-typical exemplars. Together, these findings indicate that memory transformation does not resolve the specificity–generalisation trade-off by sacrificing perceptual detail. Rather, the system simultaneously broadens the cues that can access a trace, preserves the trace’s perceptual content when retrieval succeeds, and reconstructs from category structure when retrieval fails. These results suggest that memory transformation involves co-occurring but partially dissociable changes: retrieval becomes less tied to the encoded cue, perceptual detail remains accessible, and reconstruction becomes increasingly shaped by category structure.

## Methods

### Participants

Participants were recruited online through Prolific and received monetary compensation at a rate of £6.00 per hour. Eligibility criteria required participants to be between 18 and 35 years old, to report English as both their first and primary language, to have normal or corrected-to-normal vision, to have a Prolific approval rate of at least 95%, and to have completed at least 30 previous Prolific studies. Recruitment was restricted to participants from the United Kingdom and the United States. Participants who had taken part in previous versions, pilot studies, or related studies from our laboratory using the same stimulus set were excluded from recruitment.

As reported in the pre-registration (see below), participants were tested in batches of 28. Recruitment started with the 6-hour delay group only. After each batch, we computed mean accuracy in the categorisation task per participant (collapsed across cue conditions) and ran a Bayesian one-sample t-test against chance (0.25). Once the 6-hour group reached BF₁₀ ≥ 10 confirming robust above-chance performance, recruitment proceeded sequentially across delays. For each delay, batches were evaluated following the same Bayesian approach, testing the hypothesis that more-typical cues would outperform encoded cues in the categorisation task; data collection stopped when BF₁₀ > 3 or BF₁₀ < 0.3.

Participants were excluded if they had more than 20% timeout trials in any retrieval task, a median response time below 250 ms in any task, accuracy more than 3 SD below the task-specific group mean in any retrieval task, or fewer than eight usable trials per cue condition following trial-level exclusions. Of the 223 participants recruited, 46 were excluded for exceeding the maximum percentage of timeouts (6-hour group: n = 8; 24-hour group: n = 23; 48-hour group: n = 15). No additional participants were excluded under the remaining criteria. The final sample comprised 177 participants: 56 in the 6-hour group, 64 in the 24-hour group, and 57 in the 48-hour group. Participants ranged in age from 19 to 35 years (M = 29.27, SD = 4.34). The sample was predominantly female (65.0%, n = 115), with 59 male participants (33.3%), two participants identifying as another gender (1.1%), and one preferring not to report a gender (0.6%). Most participants reported the United Kingdom as their nationality (87.0%, n = 154), followed by the United States (11.3%, n = 20), with three participants reporting Hungarian, Irish, or Pakistani nationality (0.6% each). Similarly, English was the primary reported language for almost all participants (98.3%, n = 174). The majority of participants were right-handed (85.3%, n = 151), with 25 left-handed participants (14.1%) and one ambidextrous participant (0.6%).

The study was approved by the Ethics Committee for Human Research of the University of Granada (CEIH approval code: 4788/CEIH/2025). All participants provided informed consent online before beginning the experiment.

### Stimuli

Stimuli were colour images of everyday objects drawn from the THINGS database (Hebart et al., 2019). Typicality values for each image were obtained from an independent norming study conducted by our group. The full norming dataset will be released separately. Here we describe only the relevant aspects for the present experiment.

In the norming study 539 participants (492 after exclusions) were first asked to imagine a typical exemplar of an object concept (e.g., dog) and then arranged six images from that concept according to their similarity to the imagined exemplar. Rankings were converted into a typicality index ranging from −1 to +1 and subsequently z-scored within participant. For each image, standardised scores were evaluated using a one-sample *t*-test against zero. Images with |*t*| < 0.75 were tagged as eligible encoded exemplars (average typicality), whereas images with *t* ≥ 2.5 or *t* ≤ −2.5 were considered more- and less-typical alternatives, respectively.

From the normative dataset, we selected 112 object concepts and three images per concept: one encoded exemplar, one more-typical alternative, and one less-typical alternative. The complete stimulus set therefore comprised 336 images.

We examined whether the distinction between less- and more-typical cues relative to the encoded exemplar could be attributed simply to differences in low-level visual similarity, such as colour, edges, or local shape. To this end, we combined image representations extracted from CORnet-S, a deep neural network designed to approximate hierarchical processing along the primate ventral visual stream (Kubilius et al., 2019), with representational similarity analysis (Kriegeskorte et al., 2008). Cue typicality was not reliably discriminable from distance to the encoded image in the lower and intermediate network layers (V1, V2, and V4), arguing against a low-level visual similarity confound. A modest association emerged only in the IT layer, consistent with the higher-level representational differences expected from a typicality manipulation (see Supplementary Materials for details).

The encoding set in each stimulus list comprised 48 object–object pairs (96 images), with 16 pairs assigned to each retrieval-cue condition. Within each pair, one object was designated as the retrieval cue and the other as its associate. The retrieval-cue set comprised 48 images: 16 exact repetitions of the encoded exemplars, 16 more-typical alternatives, and 16 less-typical alternatives. More- and less-typical cues depicted different exemplars of the same concept as the corresponding encoded image.

To counterbalance the experimental conditions, we first created seven lists in which each object concept appeared once as a cue and once as an associate in each of the three retrieval-cue conditions. Four alternative pairing arrangements were then generated for each list, yielding 28 stimulus lists in total. These arrangements were selected to minimise the frequency with which the same two objects were paired across lists.

### Procedure

The experiment was run online using JATOS (v3; Lange et al., 2015) and jsPsych (v7.3.4; De Leeuw et al., 2023). Participants provided informed consent, completed demographic questions, and were informed of their assigned retention interval (6, 24, or 48 hours). They then completed an instruction block explaining the associative encoding task, the three subsequent retrieval tasks, and the confidence ratings, with annotated examples provided to ensure task comprehension. Participants were instructed to memorise each pair by imagining the two objects interacting and by attending to the specific visual details of both images (Fig. 1a).

They were also informed that a retrieval cue could depict either the exact encoded image or a different exemplar from the same concept, and that they should retrieve the originally associated object despite this visual change. However, participants were discouraged from relying on purely concept-based encoding by being informed that the perceptual discrimination task would require them to identify the exact image of the associated object. Importantly, they did not know which member of each pair would later serve as the cue or the associate.

Each encoding trial began with a fixation cross presented randomly between 500 and 1,000 ms, followed by the object pair. The pair remained visible for a minimum of 2 s, after which participants could press the spacebar to continue, and for a maximum of 10 s. Participants were instructed to continue only once they felt they had sufficiently encoded the pair. Each pair was presented twice across two encoding rounds. Presentation order was randomised independently in each round. During the first round, the designated retrieval cue appeared equally often on the left and right. The positions of the two objects were reversed for every pair during the second round.

Eight shape-identification trials were interspersed throughout each encoding round. These trials were designed as attentional check trials and participants were asked to identify a circle, square, triangle, or diamond using the arrow keys, with a maximum response time of 4 s. Participants were required to achieve at least 75% accuracy in each round (≥6 of 8 correct) to avoid exclusion.

The encoding and retrieval phases were separated by a retention interval of 6, 24 or 48 hours (Fig. 1b). Each participant completed only one delay condition, but the structure of both phases was identical across delay groups. Encoding times were comparable across delay conditions, as confirmed by a one-way ANOVA, *F*(2, 174) = 0.73, *p* = .485, η² = .008. During the retrieval phase, participants received more detailed instructions about the retrieval tasks, including the specific timing of each task and trial.

The retrieval test comprised the 48 studied object pairs, presented in randomised order. Sixteen trials were assigned to each of the three cue conditions. On each trial, one object from the encoded pair was presented as a retrieval cue, and participants were asked to retrieve the object with which it had been paired during encoding. Depending on the cue condition, the cue was either the original encoded image, a more-typical exemplar, or a less-typical exemplar of the same object concept (Fig. 1c). Each trial consisted of three consecutive retrieval tasks (Fig. 1d). First, in the categorisation task, participants selected the superordinate category of the associated object from four alternatives using the arrow keys. The response pool comprised 12 superordinate categories: aquatic, birds, clothes, electronics, furniture, insects, instruments, mammals, reptiles, tools, toys, and vehicles. On each trial, the correct category was presented alongside three distractor categories randomly sampled from the remaining 11 categories. The four labels were then randomly assigned to the upper, lower, left, and right arrow response positions. Consequently, both the distractor categories and the spatial position of the correct response varied across trials. The alternatives were previewed for 1 s before the cue appeared. Participants then had up to 7 s to respond using the arrow key corresponding to their selected category. Second, participants completed the semantic recall task while the retrieval cue remained on screen. They typed the name of the associated object concept (e.g., ant, knife) into an autocomplete field, which displayed matching entries from a predefined list of valid object names. A response could be submitted only if it matched an entry in this vocabulary. Participants had up to 20 s to respond. Third, participants completed the perceptual discrimination task while the cue remained visible. Of the 48 trials, 36 were Target-present trials. On these trials, three images from the associated object concept were displayed below the cue: the exact exemplar, a more-typical exemplar, and a less-typical exemplar. Participants had to select the encoded exemplar using the left, up, or right arrow key. On the remaining 12 Target-absent trials, four per cue condition, the three alternatives were exemplars from an unstudied object concept (with one reference exemplar, one more-typical alternative, and one less-typical alternative), making “none of these” the correct response. Participants selected “none” using the down arrow. The spatial positions of the three images were counterbalanced across trials, and responses were limited to 12 s. After each of these three retrieval tasks, participants rated their confidence on a scale from 0 (“not at all”) to 100 (“extremely”) within 5 s. Confidence ratings were preceded and followed by a 500-ms fixation cross. Participants were instructed to respond as quickly and accurately as possible and to make their best guess when uncertain.

### Data Preprocessing

Reaction-time analyses excluded incorrect trials and those with latencies more than ±3 SD from a participant’s mean within each condition. For analyses involving the typicality bias index at the participant level (i.e., the coupling analyses reported below), participants were required to have contributed a minimum of five more-typical or less-typical responses on Target-present error trials. Participants who fell below this threshold were excluded from the coupling analyses only and were retained in all other analyses.

### Derived Measures

A Typicality Bias Index was computed to quantify the direction of perceptual errors. This index was computed on those trials in which participants selected either the more-typical or less-typical exemplar:

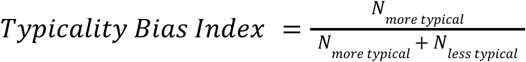

Values above .5 indicate a bias towards more-typical responses, values below .5 indicate a bias towards less-typical responses, and .5 indicates no directional bias.

We additionally derived an item-level Episodic Strength measure from performance in the semantic recall task:

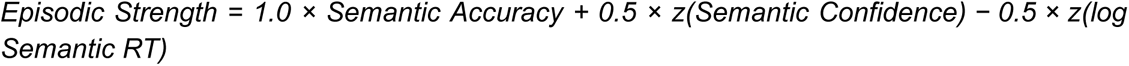

When Episodic Strength was compared across delays, confidence and RT were standardised globally to preserve between-delay differences. When it was used as a trial-level predictor of perceptual performance, these components were standardised within participants, indexing the relative strength of an item while removing stable individual differences in confidence and response speed.

### Preregistration

The design, sampling plan, and primary analyses were preregistered before data collection (https://aspredicted.org/j8493t.pdf), including the sequential Bayesian stopping rule and the exclusion criteria described above.

The primary preregistered hypothesis was that more-typical cues would support better retrieval than the originally encoded cues. This was tested with Bayesian paired-samples *t*-tests, using the Balanced Integration Score (BIS) for the categorisation task and semantic distance (the cosine dissimilarity between the typed response and the target label in THINGS embedding space) for the semantic recall task (see results in the Supplementary Material). We further preregistered mixed-design ANOVAs on accuracy, response time, BIS, and confidence, with cue condition as a within-participant factor and delay as a between-participant factor, together with planned within-delay contrasts between all cue pairs.

Two departures from the preregistration should be noted. Semantic distance is degenerate on correct trials, where it is zero by definition, so condition differences in mean semantic distance largely re-express differences in accuracy rather than indexing the graded semantic content of retrieval. We therefore based our main analyses of the semantic recall task on a composite index of retrieval strength combining accuracy, confidence, and response speed; the preregistered semantic-distance comparisons are reported in full in the Supplementary Material, as is a version of the composite incorporating semantic distance, which yields the same conclusions. The composite index was itself not preregistered, and two alternative operationalisations are reported in the Supplementary Material.

Analyses of the perceptual discrimination task were designated exploratory in the preregistration, as were all analyses relating retrieval strength and cue generalisation to perceptual accuracy and to the direction of errors. This includes the typicality bias index, the coupling analyses, and the minimum-trial threshold applied to them. These analyses are labelled as exploratory.

## Results

### Retrieval generalises to more-typical cues

First, we analysed performance relative to the memory categorisation task. Here, after presenting a cue, participants were asked to choose the category of the associated target within four options (Fig. 1d). Given that this task was the first to be answered in all retrieval trials, it allowed us to have a better estimate of the balance between accuracy and reaction times. Participants performed well across all delays and cue conditions (3×3; Fig. 2a), showing an above chance accuracy when using a series of T-tests against chance (all *p* < 0.001). A mixed ANOVA with delay (6 h, 24 h, 48 h) as a between-subject factor and cue condition (encoded, more-typical, less-typical) as a within-subject factor revealed a significant main effect of delay, *F*(2, 174) = 7.47, *p* = .0008, ηp² = .079, indicating that, although performance was robustly above chance, categorisation performance differed across retention intervals. There was also a significant main effect of cue typicality, *F*(2, 348) = 20.97, *p* < .001, ηp² = .108, as well as a significant delay × cue typicality interaction, *F*(4, 348) = 2.79, *p* = .026, ηp² = .031. Descriptively, accuracy was highest in the encoded cue condition at both 6 h (M = .535) and 24 h (M = .512), with lower performance for more-typical and less-typical cues. At 48 h, performance was lower overall, and the encoded cue and more-typical cue conditions were virtually identical (encoded: M = .372; more-typical: M = .372), while the less-typical condition remained lowest (M = .324).

**Figure 2.**
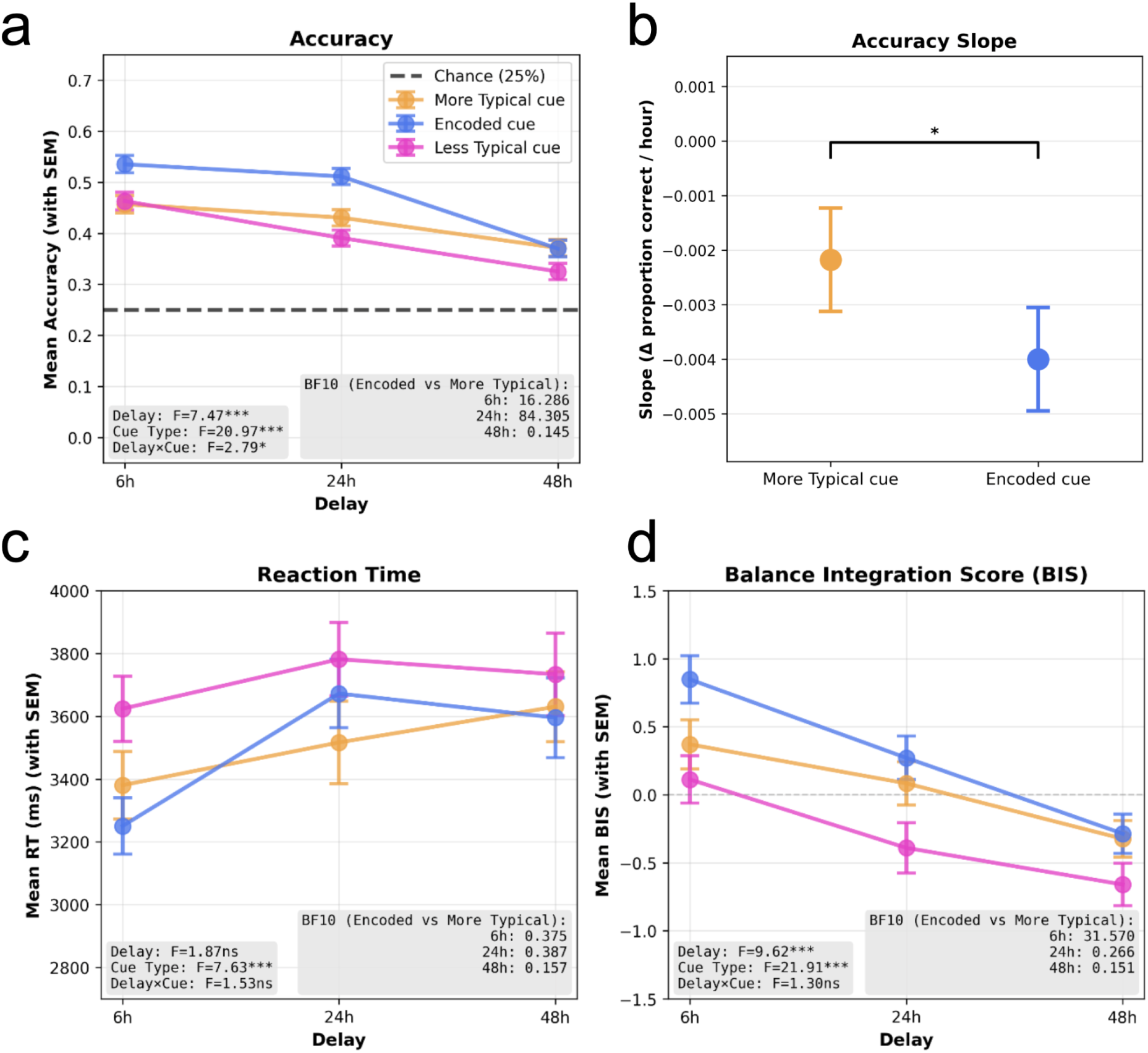
Categorisation task. Retrieval generalises to never-encoded more-typical cues by 48 h. Performance on the four-alternative superordinate-category task by retention delay (6 h, 24 h, 48 h) and cue type (encoded, More typical, Less typical). (a) Mean accuracy; dashed line indicates chance (25%). (b) Per-condition accuracy slope over delay (Δ proportion correct per hour) for encoded versus More typical cues, from a mixed-effects model; the encoded cue declines roughly twice as steeply. (c) Mean reaction time (ms) on correct trials. (d) Balanced Integration Score (BIS = z(accuracy) − z(RT)), with higher values indicating better integrated performance. Error bars denote SEM. Panels display the corresponding mixed-ANOVA terms (delay, cue typicality, delay × cue typicality) and Bayesian encoded-versus-more-typical comparisons per delay (BF₁₀). *p < .05, ***p < .001; ns, not significant.

To follow up on the interaction results, separate repeated-measures ANOVAs were conducted within each delay group. At 6 h (*n* = 56), there was a significant effect of cue typicality, *F*(2, 110) = 7.75, *p* = .0007, with accuracy highest for encoded cues (M = .535), followed by less-typical (M = .465) and more-typical cues (M = .462). At 24 h (*n* = 64), the effect of cue typicality was again significant, *F*(2, 126) = 15.68, *p* < .001, with accuracy decreasing from encoded (M = .512) to more-typical (M = .433) to less-typical cues (M = .394). At 48 h (*n* = 57), the effect of cue typicality was marginal, *F*(2, 112) = 3.00, *p* = .054; here, encoded (M = .372) and more-typical (M = .372) cues produced comparable performance, whereas less-typical cues yielded numerically lower accuracy (M = .324).

Given the a priori prediction comparing the studied image (encoded cue) against the more-typical cue, paired Bayesian *t*-tests were conducted within each delay group. For accuracy, these analyses showed evidence favouring encoded over more-typical at 6 h (Mdiff = .072, BF₁₀ = 16.29) and especially at 24 h (Mdiff = .079, BF₁₀ = 84.31). In contrast, at 48 h there was evidence for the null, with no difference between encoded and more-typical cues (Mdiff = -.0002, BF₁₀ = 0.145). Thus, categorisation accuracy was better when using studied cues (6 h, 24 h), however this benefit disappeared at longer delays (48 h), when a never-seen but more typical version of the studied trigger showed a similar performance.

To test whether accuracy declined at different rates for encoded versus more-typical cues, we fit a mixed-effects model regressing accuracy (proportion correct) on delay hours (hours, mean-centred) and cue typicality, with a random intercept per participant. The critical term was the delay hours x cue typicality interaction, which tests whether the slope of accuracy over delay differs between the two conditions. This interaction was significant (b = -0.0018 per hour, SE = 0.00073, z = -2.49, p = .013), indicating that the two conditions declined at different rates. The more-typical cue showed a shallow but significant decline (slope = -0.0022/hour, SE = 0.00095, p = .022), whereas the encoded cue declined roughly twice as steeply (slope = -0.0040/hour, SE = 0.00095; Fig. 2b). In other words, accuracy for items retrieved via their originally encoded cue eroded significantly faster than for items retrieved via a more-typical cue.

The same ANOVA on reaction times (RT; Fig. 2c) showed no main effect of delay, *F*(2, 174) = 1.87, *p* = .158, ηp² = .021, and no delay × cue typicality interaction, *F*(4, 348) = 1.53, *p* = .194, ηp² = .017. However, there was a significant main effect of cue typicality, *F*(2, 348) = 7.63, *p* = .0006, ηp² = .042, indicating that response speed varied across cue types.

Within-delay analyses showed a significant effect of cue typicality at 6 h, *F*(2, 110) = 8.99, *p* = .0002, with participants responding fastest to encoded cues (M = 3250.5 ms), more slowly to more-typical cues (M = 3380.5 ms), and slowest to less-typical cues (M = 3624.0 ms). At 24 h, the effect was not significant, *F*(2, 126) = 2.29, *p* = .106, although descriptively responses were fastest for more-typical cues (M = 3516.6 ms), followed by encoded (M = 3672.5 ms) and less-typical cues (M = 3781.8 ms). At 48 h, there was also no significant effect of cue typicality, *F*(2, 112) = 1.32, *p* = .270; mean RTs were 3595.6 ms for encoded, 3630.5 ms for more-typical, and 3733.7 ms for less-typical cues.

Bayesian planned comparisons between encoded and more-typical cues did not provide evidence for a reliable RT advantage for encoded cues at any delay. At 6 h, the evidence was inconclusive to weakly against an encoded-cue speed advantage (Mdiff = -130.1 ms; BF₁₀ = 0.375). The same pattern held at 24 h (Mdiff = 155.9 ms; BF₁₀ = 0.387) and 48 h (Mdiff = -35.0 ms; BF₁₀ = 0.157), with the latter two comparisons providing evidence for the null.

To assess overall performance while accounting for speed–accuracy trade-offs, we analysed the Balanced Integration Score (BIS; Liesefeld et al., 2015; Liesefeld & Janczyk, 2019; Fig. 2d), computed as z(accuracy) − z(RT), such that higher values indicate better overall performance. The mixed ANOVA on BIS revealed a significant main effect of delay, F(2, 174) = 9.62, p < .001, ηp² = .100, showing that integrated performance declined across longer delays. There was also a significant main effect of cue typicality, F(2, 348) = 21.91, p < .001, ηp² = .112, but no delay × cue typicality interaction, F(4, 348) = 1.30, p = .268, ηp² = .015.

Descriptively, BIS was highest at 6 h, intermediate at 24 h, and lowest at 48 h, consistent with declining performance over time. Across delays, encoded cues tended to show the highest BIS values, whereas less-typical cues consistently produced the lowest values. Within-delay analyses confirmed a significant effect of cue typicality at 6 h, F(2, 110) = 15.14, p < .001, with BIS highest for encoded cues (M = 0.848), followed by more-typical (M = 0.371) and less-typical cues (M = 0.113). A similar effect emerged at 24 h, F(2, 126) = 7.22, p = .001, with BIS values of 0.272, 0.084, and -0.390 for encoded, more-typical, and less-typical cues, respectively. At 48 h, the cue typicality effect remained significant, F(2, 112) = 4.23, p = .017, driven mainly by poorer performance for less-typical cues (M = -0.660) relative to encoded (M = -0.286) and more-typical cues (M = -0.324), which were again very similar.

Bayesian encoded-versus-more-typical comparisons on BIS further confirmed this pattern. At 6 h, there was clear evidence favouring encoded over more-typical (Mdiff = 0.478, BF₁₀ = 31.57). At 24 h, however, the data supported the null rather than an encoded cue advantage (Mdiff = 0.187, BF₁₀ = 0.266). Likewise, at 48 h, there was evidence for no difference between encoded and novel but more-typical cues (Mdiff = 0.038, BF₁₀ = 0.151).

Overall, the categorisation task showed that performance declined with increasing delay, particularly in accuracy and BIS. Across measures, less typical versions of studied cues were generally associated with the poorest performance. The categorisation results reveal a theoretically important shift across delay: whereas encoded cues produced better accuracy than more-typical cues at 6 h and 24 h, this “encoding” advantage disappeared by 48 h. At 48 h the more-typical cue matched the retrieval performance of the originally encoded cue, despite not being the studied stimulus itself. This pattern suggests that delayed retrieval can be supported by a more typical representation, consistent with the idea that the functional match between cue and memory changes over time. Importantly, this cue-generalisation towards more typical exemplars is difficult to reconcile with a simple verbal-label account of the association. Although both the more-typical and the less-typical cues share the same verbal label as the encoded cue, only the more-typical cues converged on encoded-cue performance by 48 h: the encoded-vs-more-typical comparison yielded evidence for the null on both accuracy (BF₁₀ = 0.145) and BIS (BF₁₀ = 0.151), whereas the corresponding encoded-versus-less-typical comparisons did not provide evidence for convergence (accuracy BF₁₀ = 1.07; BIS BF₁₀ = 2.75), with less-typical cues remaining numerically lowest. A purely verbal-label account would predict both label-matched cues to behave alike; instead, the clearest evidence for convergence was observed for the more-typical exemplar.

Two features argue that this convergence is not merely a floor effect. First, it is a difference in rate, not only level: the encoded cues’ accuracy declined roughly twice as fast as the more-typical cues’ (interaction b = −0.0018/hour, p = .013), so the conditions met because the encoded advantage was actively eroding, not because performance was compressed into a shared floor. Second, convergence was selective. A generic floor would draw all three cue types together, yet the less-typical cue did not match the encoded cue on both accuracy and BIS, whereas only the more-typical cue reached convergence (BF₁₀ = 0.145 and 0.151). Converging performance also remained well above chance (.37 vs .25).

These results are difficult to reconcile with a static encoding-specificity view, which would predict a persistent advantage for the originally studied image. Instead, the convergence between encoded and more-typical cues at 48 h suggests that the cue–trace match is non-stationary: as memories transform, a never-seen but more typical exemplar becomes as effective a retrieval probe as the originally encoded one. Such patterns are consistent with the Dynamic-Cueing hypothesis and its core claim that the most effective cue depends on the current representational state of the trace, which evolves over consolidation.

### Perceptual details remain accessible and cue-dependent

In the perceptual discrimination task, participants selected the target among four response options (the encoded, a more-typical, a less-typical variant of the target, plus a “none of these” option), with 25% of trials being lure trials in which all options were replaced by a novel concept, with images having the same three different levels of typicality. Thus, chance performance in this task was 25%.

Accuracy was analysed using a mixed ANOVA with delay as a between-subject factor and cue typicality as a within-subject factor (Fig. 3a). The analysis revealed no main effect of delay, *F*(2, 174) = 1.51, *p* = .225, ηp² = .017, but a significant main effect of cue typicality, *F*(2, 348) = 3.75, *p* = .024, ηp² = .021. The delay × cue typicality interaction was not significant, *F*(4, 348) = 0.86, *p* = .489, ηp² = .010. Collapsed across delays, accuracy was highest for encoded cues (*M*= .707, *SD* = .199), followed by more-typical (*M* = .682, *SD* = .205) and less-typical cues (*M* = .677, *SD* = .213). Descriptively, performance was relatively similar across cue types at 6 h (encoded: *M* = .723, more-typical: *M* = .696, less-typical: *M* = .711) and 48 h (encoded: *M* = .668, more-typical: *M* = .662, less-typical: *M* = .632), whereas at 24 h encoded cues showed better performance (encoded: *M* = .727, more-typical: *M* = .686, less-typical: *M* = .687).

**Figure 3.**
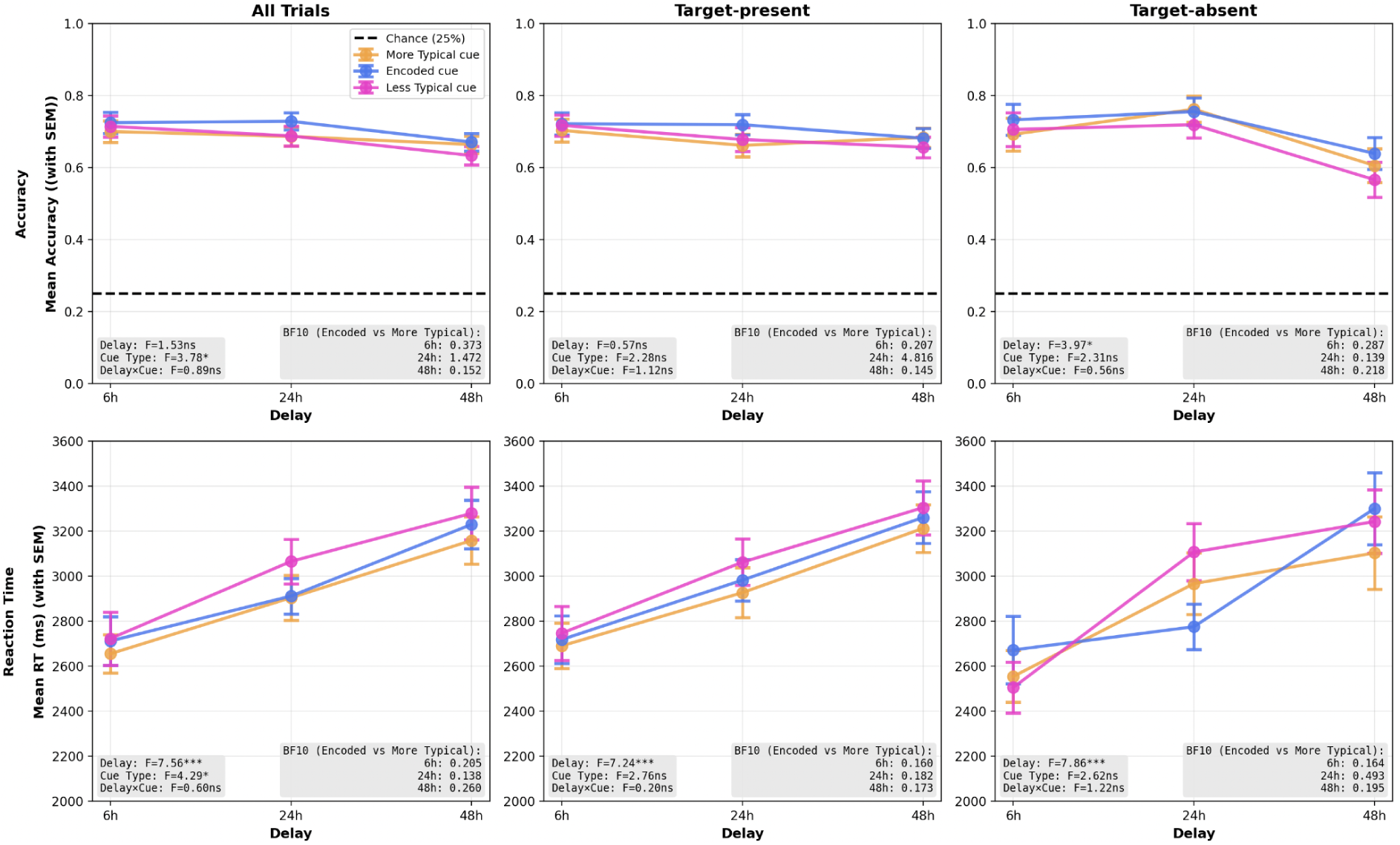
Perceptual discrimination task: concrete detail remains accessible but cue-dependent. Exemplar-recognition performance by retention delay and cue type, shown separately for all trials, Target-present trials, Target-absent trials. (**a**) Mean accuracy; the dashed line marks chance (25%). Accuracy remained above chance in every delay × cue-type combination and did not decline systematically with delay; only rejection of novel-concept lures worsened at 48 h. (**b**) Mean reaction time on correct trials, which slowed reliably with delay and was graded by cue typicality (fastest for more-typical, slowest for less-typical cues). Error bars denote ± SEM. In-panel annotations report the mixed-ANOVA effects and the Bayesian comparison of encoded versus more-typical cues.

Follow-up paired comparisons within each delay group showed that at 24 h, encoded cues yielded higher accuracy than both more-typical cues, *t*(63) = -2.20, *p* = .031, *d* = -0.204, and less-typical cues, *t*(63) = 2.09, *p* = .041, *d* = 0.194. No other within-delay cue-typicality contrasts reached significance (all *ps* ≥ .095). Bayesian comparisons between encoded and more-typical cues provided inconclusive evidence at 6 h (BF₁₀=0.373), anecdotal evidence for a difference at 24 h (BF₁₀ = 1.472), and moderate evidence favouring the null at 48 h (BF₁₀ = 0.152). Thus, although accuracy showed a modest overall advantage for encoded cues, there was no reliable evidence that this advantage changed systematically across delay, and by 48 h the data supported comparable accuracy for encoded and more-typical cues.

Accuracy on Target-absent trials was analysed separately using the same mixed ANOVA structure. This analysis revealed a significant main effect of delay, *F*(2, 174) = 3.97, *p* = .021, ηp² = .044, but no main effect of cue typicality, *F*(2, 348) = 2.31, *p* = .101, ηp² = .013, and no delay × cue typicality interaction, *F*(4, 348) = 0.56, *p* = .695, ηp² = .006. Collapsed across cue types, new-concept performance was highest at 24 h (*M* = .745, *SD* = .294), intermediate at 6 h (*M*= .710, *SD* = .335), and lowest at 48 h (*M* = .603, *SD* = .352). Pairwise comparisons across delays showed that performance at 24 h was higher than at 48 h for all three cue conditions: more-typical, *t*(119) = 2.67, *p* = .0086, *d* = 0.484; encoded, *t*(119) = 2.02, *p* = .046, *d* = 0.367; and less-typical, *t*(119) = 2.53, *p* = .0128, *d* = 0.457. No within-delay comparisons between cue types were significant (all *ps* ≥ .213). Consistent with this, Bayesian comparisons between encoded and more-typical cues favoured the null at all three delays (BF₁₀ = 0.287, 0.139, and 0.218 for 6 h, 24 h, and 48 h, respectively). Therefore, the main pattern for Target-absent trials was a delay-related reduction in performance at 48 h rather than a reliable effect of cue typicality.

Accuracy on Target-present trials, in which the encoded target was presented alongside a more-typical and a less-typical alternative from the same concept, was also analysed separately. In contrast to the Target-absent analysis, this ANOVA showed no main effect of delay, *F*(2, 174) = 0.57, *p* = .566, ηp² = .007, no main effect of cue typicality, *F*(2, 348) = 2.28, *p* = .104, ηp² = .013, and no delay × cue typicality interaction, *F*(4, 348) = 1.12, *p* = .346, ηp² = .013. Descriptively, Target-present accuracy remained fairly stable across delays and cue types. At 24 h, however, an exploratory within-delay comparison showed higher performance for encoded than more-typical cues, *t*(63) = -2.80, *p*= .0067, *d* = -0.242, with moderate Bayesian evidence for a difference (BF₁₀ = 4.816). In contrast, the corresponding Bayesian comparisons favoured the null at 6 h (BF₁₀ = 0.207) and 48 h (BF₁₀ = 0.145). No other pairwise comparisons were significant (all *ps* ≥ .078). Given the absence of any significant omnibus effect and the lack of evidence for an encoded-versus-more-typical difference at the other delays, this isolated 24 h contrast should be interpreted cautiously.

Importantly, performance remained robustly above chance (.25) in every delay × cue typicality condition, both overall and when Target-present and Target-absent trials were considered separately (all ps < .001). Thus, even where encoded and more-typical cues produced comparable performance, this occurred well above chance.

Reaction times on correct trials were analysed with the same mixed ANOVA, after applying the preregistered RT exclusions (Fig. 3b). Unlike accuracy, RT was sensitive to delay. Responses slowed progressively across retention intervals, *F*(2, 174) = 7.56, *p* < .001, ηp² = .080 (6 h: *M* = 2697 ms; 24 h: *M* = 2961 ms; 48 h: *M* = 3223 ms). RTs also varied with cue typicality, *F*(2, 348) = 4.29, *p* = .014, ηp² = .024, with no delay × cue typicality interaction, *F*(4, 348) = 0.60, *p* = .663, ηp² = .007. Collapsed across delay, participants were fastest for more-typical cues (*M* = 2908 ms), slowest for less-typical cues (*M* = 3027 ms), and intermediate for encoded cues (*M* = 2952 ms). Pairwise comparisons suggested that this effect was clearest at 24 h, where less-typical cues were slower than both more-typical cues, *t*(63) = −2.31, *p* = .024, *d* = −0.203, and encoded cues, *t*(63) = −2.53, *p* = .014, *d* = −0.216. No other within-delay comparisons reached significance. Importantly, the encoded cue conferred no reliable speed advantage over the never-seen more-typical cue. Bayesian paired comparisons between encoded and more-typical cues favoured the null at all three delays, BF₁₀=0.205, 0.138, and 0.260 for 6 h, 24 h, and 48 h, respectively. Thus, although responses became slower with delay and were numerically fastest for more-typical cues, there was no evidence that the originally studied cue provided a privileged speed benefit in the perceptual discrimination task.

Separating trial types showed that the delay-related slowing was general, emerging for both Target-absent trials, *F*(2, 144) = 7.86, *p* < .001, ηp² = .098, and Target-present trials, *F*(2, 170) = 7.24, *p* < .001, ηp² = .078. In contrast, the cue typicality effect was weaker when trial types were analysed separately, falling short of significance for both new-concept trials, *F*(2, 288) = 2.62, *p* = .075, ηp² = .018, and associated-item trials, *F*(2, 340) = 2.76, *p* = .065, ηp² = .016. Neither analysis showed a delay × cue typicality interaction, both *ps* ≥ .303. Bayesian comparisons likewise provided little evidence for an encoded-versus-more-typical RT difference. For Target-present trials, the data favoured the null at all three delays (BF₁₀ = 0.160, 0.182, and 0.173), while for Target-absent trials the evidence favoured the null at 6 h and 48 h (BF₁₀ = 0.164 and 0.195) and was inconclusive at 24 h (BF₁₀=0.493). Thus, the clearest RT effect when separating trial types was a robust slowing with delay, whereas cue-related differences in response speed were modest and less stable across trial-type subsets.

Overall, accuracy in the perceptual discrimination task did not decline systematically across delay; only Target-absent performance showed a delay effect, with worse lure rejection at 48 h, whereas Target-present performance showed no reliable omnibus effects, and performance remained robustly above chance in every condition. Response times, by contrast, slowed reliably with delay. Cue typicality had an effect on RT, driven mainly by poorer performance for less-typical cues, but Bayesian comparisons provided no evidence for a reliable encoded-versus-more-typical speed advantage. Together, these results suggest that concrete perceptual information remained accessible across delays, while retrieval became slower and less efficient over time.

### Errors reveal a typicality drift

In the perceptual discrimination task, the three image alternatives represented three typicality levels relative to a given concept (see Methods). On Target-present trials, these were the encoded target (the correct response), a more-typical alternative, and a less-typical alternative. On Target-absent trials, the three images belonged to a never-encoded concept, and “none of these” was the correct response (Fig. 4a).

**Figure 4.**
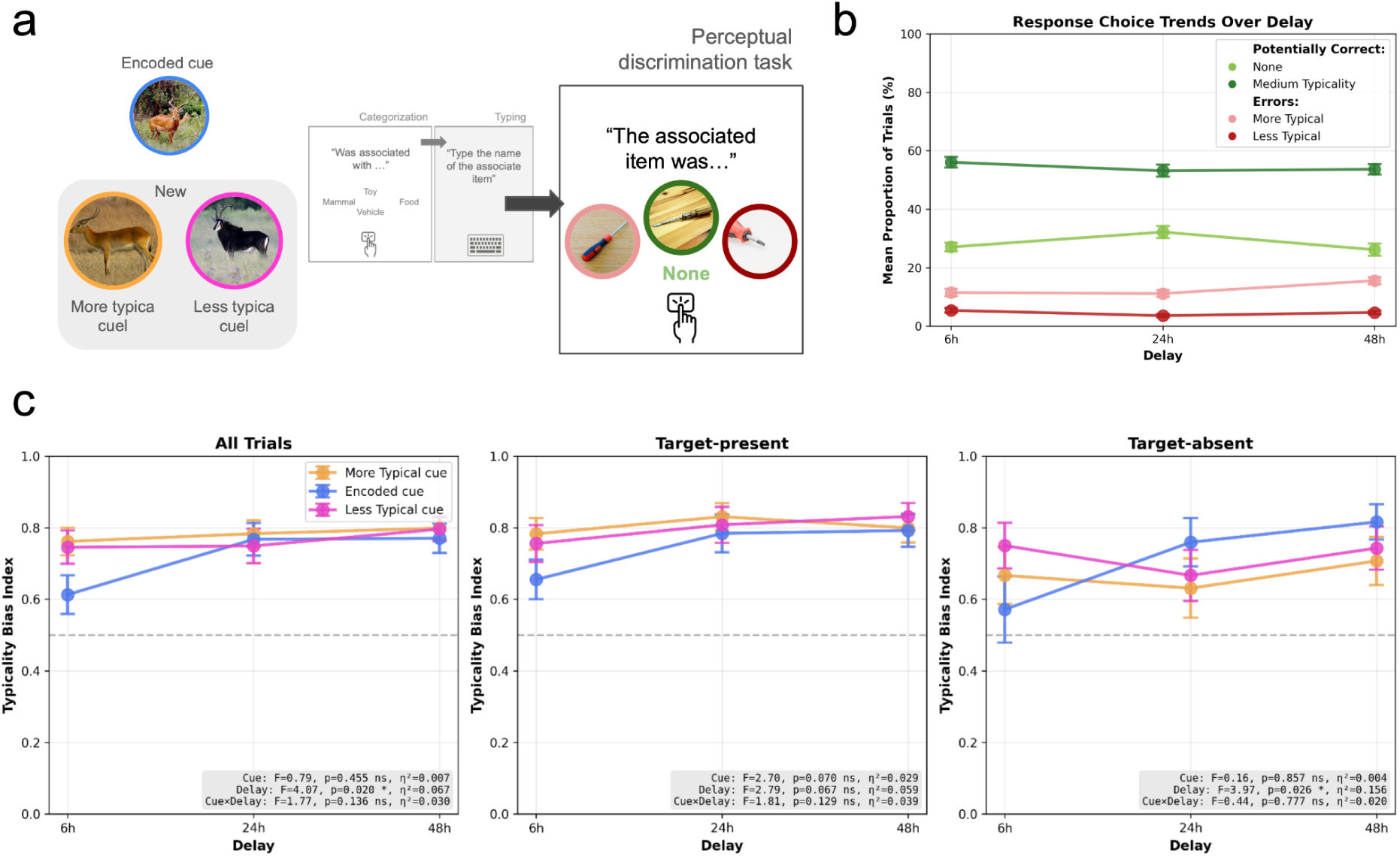
Perceptual errors become increasingly biased toward category-typical exemplars. (**a**) Trial schema for the perceptual discrimination task. After the categorisation and semantic recall tasks, participants selected the associated object among the exact encoded exemplar (reference typicality; the target), a more-typical lure and a less-typical lure of the same concept, or “none” when all three options came from a novel concept. (**b**) Mean proportion of trials assigned to each response category across delay: correct selection of the target (reference/medium typicality), correct “none” responses, and the two error types (More Typical, Less Typical). More-typical errors increased with delay while less-typical errors remained rare, so that by 48 h more-typical errors outnumbered less-typical errors roughly 3:1. (**c**) Typicality Bias Index computed per participant over lure-selection errors, by delay and cue type, for all, target-present, and target-absent errors; the dashed line at 0.5 indicates no directional bias. The index exceeded 0.5 at every delay and increased with retention intervals. Error bars denote ± SEM; in-panel annotations report the mixed-ANOVA effects (cue typicality, delay, cue typicality × delay).

Across all trials and delays, more-typical and less-typical choices showed clearly divergent profiles across delays (Fig. 4b). More-typical responses were consistently the more frequent of the two error types and grew with delay, rising from M = 11.5% (SEM = 1.32) at 6h to M = 15.5% (SEM = 1.22) at 48h. Less-typical responses, in contrast, were rare throughout and became rarer still, falling from M = 5.4% (SEM = 0.80) at 6h to M = 3.6% (SEM = 0.55) at 24h and M = 4.7% (SEM = 0.64) at 48h. By 48h, more-typical choices outnumbered less-typical choices around 3-to-1, indicating that as delay increased, errors were increasingly pulled toward more-typical alternatives rather than toward less-typical ones.

Participant-level analyses confirmed this descriptive pattern. For each delay group, we compared the proportion of more- and less-typical responses among more/less-typical errors using paired-samples t-tests. At 6 h, more-typical responses were more frequent than less-typical responses, M = 71.3% versus 28.7%, *t*(48) = 5.619, *p* < .001, Cohen’s *d* = 0.811. The same pattern was observed at 24 h, *M* = 76.8% versus 23.2%, *t*(56) = 10.374, *p* < .001, *d* = 1.386, and at 48 h, *M* = 79.3% versus 20.7%, *t*(56) = 11.385, *p* < .001, *d* = 1.521. Collapsing across delays, *more-typical* responses were substantially more frequent than *less-typical* responses, *M* = 76.0% versus 24.0%, *t*(162) = 15.143, *p* < .001, *d* = 1.190.

We then repeated the same analyses separately for Target-absent and Target-present error trials. For Target-present errors, the bias toward more-typical responses was reliable and numerically stronger. At 6 h, more-typical responses were more frequent than less-typical responses, *M* = 73.5% versus 26.5%, *t*(48) = 5.919, *p* < .001, *d* = 0.854. At 24 h, more-typical responses again exceeded less-typical responses, *M* = 79.8% versus 20.2%, *t*(54) = 9.136, *p* < .001, *d* = 1.243. At 48 h, this pattern remained highly reliable, *M* = 81.3% versus 18.7%, *t*(53) = 12.509, *p* < .001, *d* = 1.718. Collapsing across delays, Target-present errors showed a strong typicality bias, *M* = 78.3% versus 21.7%, *t*(157) = 15.029, *p* < .001, *d* = 1.199.

For Target-absent errors, the typicality bias was reliable at all delays. At 6 h, more-typical responses exceeded less-typical responses, *M* = 67.4% versus 32.6%, *t*(35) = 3.034, *p* = .0045, *d* = 0.513. At 24 h, the same pattern was observed, *M* = 70.4% versus 29.6%, *t*(44) = 4.737, *p*< .001, *d* = 0.714. At 48 h, more-typical responses again exceeded less-typical responses, *M* = 71.7% versus 28.3%, *t*(48) = 4.649, *p* < .001, *d* = 0.671. Collapsed across delays, the Target-absent error analysis showed a reliable typicality bias, *M* = 70.1% versus 29.9%, *t*(129) = 7.211, *p* < .001, *d* = 0.635.

We next examined whether errors in the perceptual discrimination task were biased towards more gist-like, typical representations. To formalise this pattern into a single measure per participant, we computed a Typicality Bias Index (see Methods), restricted to trials on which participants chose either the more- or the less-typical exemplar: Values above .5 indicate a bias toward more typical responses, values below .5 indicate a bias toward less typical responses, and .5 indicates no directional bias.

We tested whether the Typicality Bias Index varied as a function of delay and cue typicality using mixed ANOVAs, with delay as a between-subject factor and cue typicality as a within-subject factor (Fig. 4c). For all error trials, there was a significant main effect of delay, *F*(2, 114) = 4.074, *p* = .0196, η²p = .0667. The main effect of cue typicality *F*(2, 228) = 0.791, *p* = .4546, η²p = .0069, and the cue typicality × delay interaction was not significant, *F*(4, 228) = 1.767, *p* = .1364, η²p = .0301.

For Target-present errors, we did not find a main effect of delay, *F*(2, 89) = 2.790, *p* = .0668, η²p = .0590, or main effect of cue typicality, *F*(2, 178) = 2.702, *p* = .0698, η²p = .0295, and the cue typicality x delay interaction was not significant, *F*(4, 178) = 1.808, *p* = .1293, η²p = .0390. However, for Target-absent errors, the main effect of delay was also significant, *F*(2, 43) = 3.972, *p* = .0261, η²p = .1559. Neither the main effect of cue typicality, *F*(2, 86) = 0.155, *p* = .8566, η²p = .0036, nor the cue typicality x delay interaction, *F*(4, 86) = 0.444, *p* = .7765, η²p = .0202, was significant.

Together, these analyses show that errors in the perceptual discrimination task were systematically biased towards more-typical rather than less-typical target exemplars. This typicality bias was observed at every delay and for both target-absent and target-present errors, suggesting that when participants failed to retrieve the precise target representation, their responses tended to drift towards more prototypical category information. The delay effects indicate some modulation of the magnitude of this bias over time, particularly in the full error set and in target-absent errors. However, because the cue typicality x delay interactions were not significant, there was no clear evidence that this drift-like bias depended on the typicality of the retrieval cue over time.

### Episodic strength predicts perceptual accuracy but not bias

We next tested whether episodic memory strength modulated performance in the perceptual discrimination task. Specifically, we asked whether stronger episodic memory predicted better perceptual accuracy, and whether the bias towards more-typical responses among perceptual errors could be explained by weak episodic memory.

Episodic strength was estimated from semantic-cued recall performance for the same item (see Methods; Fig. 5a). Higher values therefore indicated more accurate, more confident, and faster semantic retrieval. Because episodic strength served two distinct analytical purposes, the continuous components were standardised differently for the two analyses. When episodic strength was treated as an outcome and compared across delays, confidence and RT were standardised globally so that between-delay differences were preserved. When episodic strength was used as a trial-level predictor of perceptual performance, confidence and RT were standardised within participants, capturing the strength of each item relative to that participant’s own performance and removing stable individual differences in confidence and response speed.

**Figure 5.**
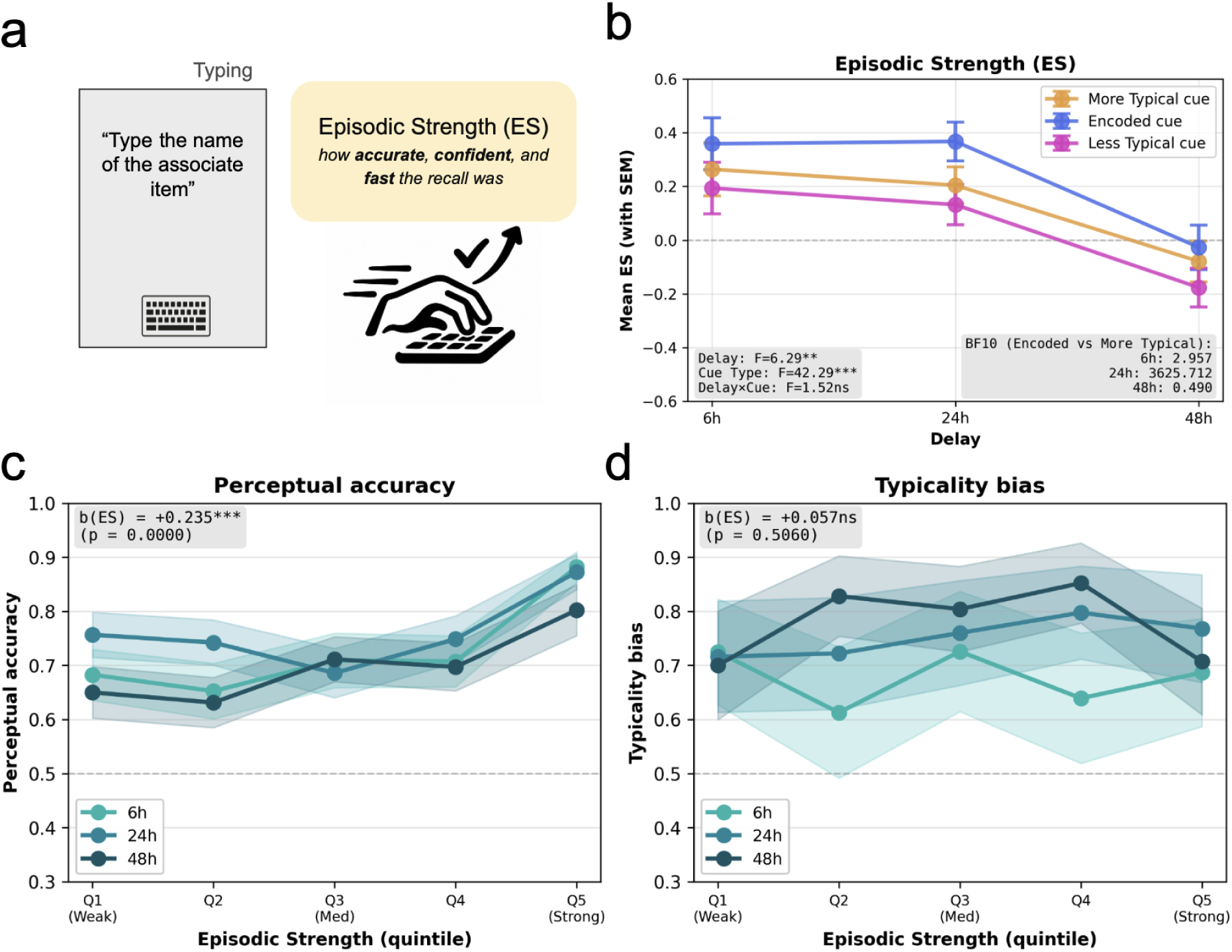
Episodic strength predicts perceptual accuracy but not the direction of errors. (**a**) Per-item episodic strength (ES) was derived from semantic-cued recall indexing accurate, confident and fast retrieval. (**b**) Mean ES by delay and cue type; error bars denote ± SEM, and in-panel annotations report the mixed-ANOVA effects and Bayesian encoded-versus-more-typical comparisons (BF₁₀). ES was comparable at 6 h and 24 h and lower at 48 h. (**c**) Perceptual accuracy as a function of within-participant episodic-strength quintile (Q1, weakest; Q5, strongest), plotted separately by delay; stronger episodic memory predicted higher perceptual accuracy (logistic GEE), with no episodic-strength × delay interaction. (**d**) Typicality Bias Index by episodic-strength quintile and delay; episodic strength did not predict error direction, whereas typicality bias increased with delay. Shaded areas denote 95% confidence intervals; in-panel annotations report the episodic-strength coefficient from the GEE model.

We first characterised the accuracy-based episodic-strength measure across experimental conditions (Fig. 5b). A mixed ANOVA revealed significant main effects of delay, *F*(2, 174) = 6.29, *p* = .002, ηp² = .067, and cue typicality, *F*(2, 348) = 42.29, *p* < .001, ηp² = .196. Their interaction was not significant, *F*(4, 348) = 1.52, *p* = .197, ηp² = .017. Collapsed across cue conditions, episodic strength was comparable after 6 h (*M* = 0.272, *SD* = 0.723) and 24 h (*M* = 0.235, *SD* = 0.582), but lower after 48 h (*M* = −0.094, *SD* = 0.583).

Planned Bayesian comparisons further tested whether episodic strength differed between the originally encoded image and the more-typical exemplar. Evidence for this difference was limited at 6 h (encoded: *M* = 0.359; more-typical: *M* = 0.264; BF₁₀ = 2.96), but extremely strong at 24 h (encoded: *M* = 0.367; more-typical: *M* = 0.205; BF₁₀ = 3625.71). At 48 h, the evidence was inconclusive and slightly favoured the null hypothesis (encoded: *M* = −0.027; more-typical: *M* = −0.079; BF₁₀ = 0.49). Thus, the advantage in episodic strength for the originally studied cue was most clearly expressed at the 24h delay and did not increase over time. The selective advantage for more-typical exemplars over time was not shared by the less-typical cue. Although the encoded cue produced stronger episodic strength than the less-typical cue at both delays (6 h: M = 0.359 vs 0.194, BF₁₀ = 2685.54; 48 h: M = −0.027 vs −0.176, BF₁₀ = 120.09), the more-typical and less-typical cues were statistically indistinguishable at 6 h (M = 0.264 vs 0.194; BF₁₀ = 0.64) yet diverged by 48 h, with the more-typical cue now yielding reliably higher episodic strength (M = −0.079 vs −0.176; BF₁₀ = 15.37). Thus the emerging episodic-strength benefit for more-typical exemplars over delay was specific to that cue and did not extend to the less-typical exemplar, even though both share the same verbal category label.

For perceptual accuracy, analyses were restricted to target-present trials. Of 6,073 target-present perceptual trials, 5,825 could be matched to semantic recall task data and were included. The final dataset included 176 participants, with 4,258 correct responses and 1,567 errors, corresponding to 73.1% accuracy. Accuracy was 73.9% at 6 h, 76.1% at 24 h, and 69.2% at 48 h. Episodic strength had a mean of 0.145, SD = 0.891.

A logistic GEE model with exchangeable within-participant correlation showed that episodic strength significantly predicted perceptual accuracy, β = 0.235, SE = 0.032, z = 7.24, p < .001, 95% CI [0.171, 0.298]. The episodic strength × delay interaction was not significant, Wald χ²(2) = 1.96, p = .375. Thus, stronger item-level memory predicted better perceptual discrimination, and this relationship did not differ reliably across delays (Fig. 5c).

We then tested whether episodic strength explained the direction of perceptual errors. To isolate typicality drift, the analysis was restricted to errors in which participants selected either the more-typical or the less-typical exemplar.

A logistic GEE model tested whether these errors were more likely to be directed towards the more-typical exemplar as a function of episodic strength and delay (Fig. 5d). Episodic strength did not predict typicality bias, β = 0.058, SE = 0.086, z = 0.67, p = .506, 95% CI [-0.112, 0.227]. In contrast, delay significantly predicted typicality bias. Relative to 6 h, more-typical errors were more likely at 24 h, β = 0.376, SE = 0.180, z = 2.10, p = .036, 95% CI [0.025, 0.728], and at 48 h, β = 0.534, SE = 0.186, z = 2.88, p = .004, 95% CI [0.170, 0.898].

An interaction model showed no evidence that the association between episodic strength and typicality bias varied by delay. The episodic strength × delay interaction was not significant, Wald χ²(2) = 1.98, p = .372. Altogether, these analyses indicate that episodic strength predicted whether participants could perceptually identify the target, but not whether their errors were pulled towards the more-typical exemplar. Thus, weak item-level memory contributed to perceptual failure in general, but did not explain the direction of the error once a perceptual error occurred.

These results dissociate two aspects of performance in the perceptual discrimination task. Episodic strength predicted whether participants correctly identified the target, suggesting that stronger item memory supported perceptual accuracy. However, among more-/less-typical errors, episodic strength did not predict whether participants selected the more-typical exemplar. Instead, typicality bias increased with retention delay. This pattern suggests that weak episodic memory contributes to perceptual failure in general, but does not explain the direction of the error once a perceptual error occurs. The drift towards typical exemplars is therefore more consistent with an increasing influence of category structure over time than with a mere weakening of episodic memory. A similar pattern of results was obtained across two other approaches to estimate episodic strength (see Supplementary Materials).

### Perceptual typicality drift and cue generalisation are not reliably coupled

The preceding analyses showed that two transformation signatures emerged with consolidation delay. Errors in the perceptual discrimination task drifted towards more-typical concept exemplars, and previously-unstudied more-typical cues became as effective as the encoded cues at 48 hours. An important question is whether these two signatures reflect a single underlying transformation of the memory representation, in which case their magnitudes should covary across individuals, or whether they reflect partially independent processes that each scale with delay through separate mechanisms.

We tested this question in participants with a minimum of five more-typical or less-typical responses on target-present trials, which provided a stable estimate of the bias index (total n = 121; 6h n = 33, 24h n = 42, 48h n = 46). In a multiple regression predicting the typicality bias index from cue generalisation (more-typical cue accuracy - encoded cue accuracy) while controlling for delay group, the coupling coefficient was small and not significant (Fig. 6a; β = -0.022, t(116) = -0.28, p = 0.78, 95% CI [-0.176, +0.133]). The 95% confidence interval rules out coupling effects larger than about |β| = 0.18 in this sample. Adding overall semantic recall accuracy as a covariate to address the possibility that shared dependence on memory strength was masking a relationship produced essentially the same estimate (β = -0.022, p = 0.78), and the cue-generalisation × delay interaction did not improve model fit (F(2, 114) = 1.29, p = 0.28), indicating that the absence of coupling did not vary across delays.

**Figure 6.**
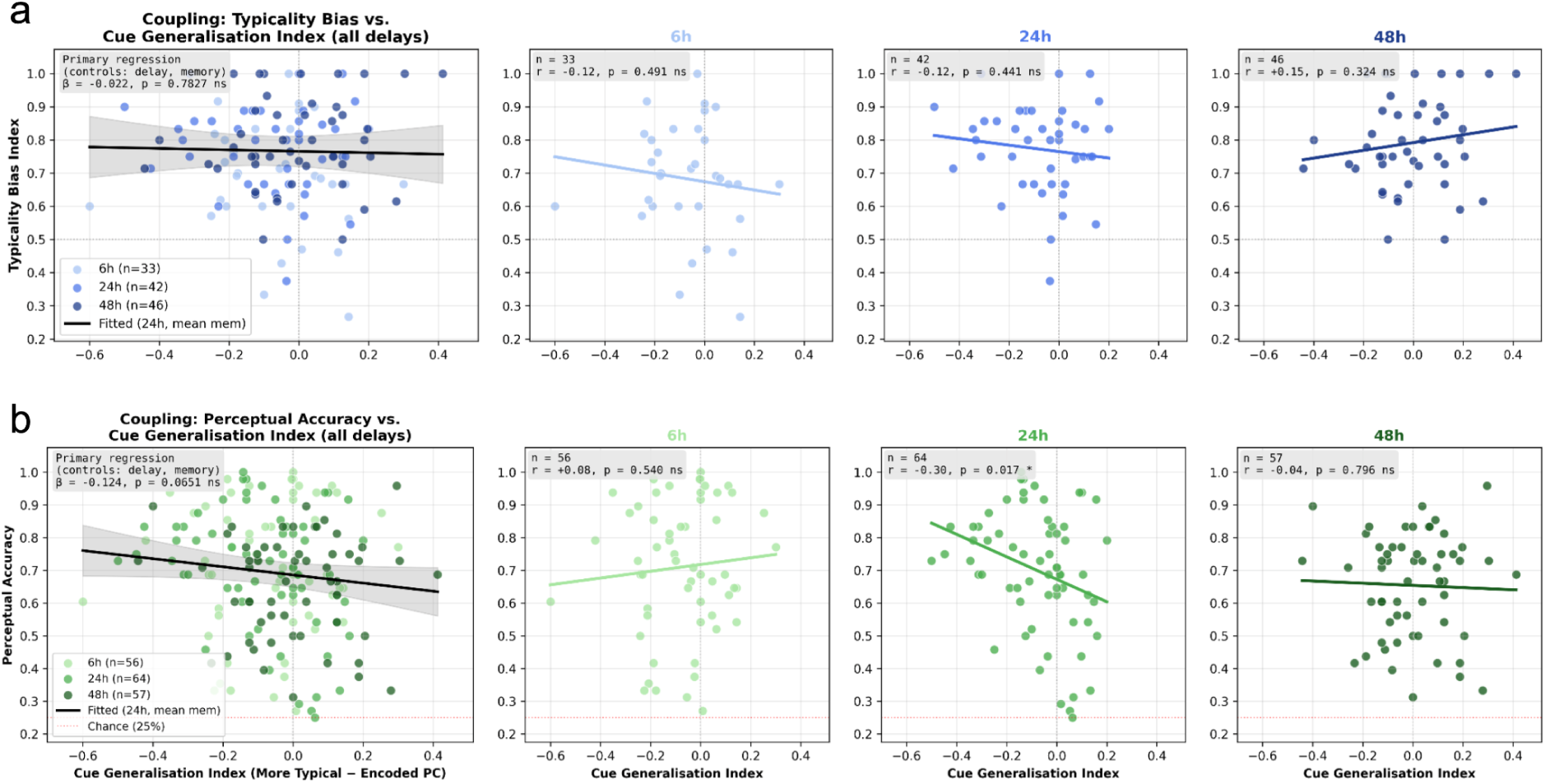
Typicality drift and cue generalisation show no reliable overall coupling. Relationship between the two transformation signatures, pooled across delays (leftmost panel, with the fitted regression controlling for delay group) and within each delay group (6 h, 24 h, 48 h). Each point is one participant. (**a**) Typicality Bias Index against the cue-generalisation index (more-typical minus encoded-cue categorisation accuracy); the association was small and non-significant overall and within every delay. Analyses were restricted to participants contributing ≥ 5 more-typical or less-typical lure selections (total *n* = 121; 6 h *n* = 33, 24 h *n* = 42, 48 h *n* = 46). (**b**) Perceptual accuracy (all trials) against the cue-generalisation index (full sample, *n* = 177; 6 h *n* = 56, 24 h *n* = 64, 48 h *n* = 57); the dashed line marks chance (25%). The overall association was marginal and negative, driven almost entirely by a significant negative correlation at 24 h and absent at 6 h and 48 h. Shaded regions denote 95% confidence intervals around the fitted lines. \**p* < .05; ns, not significant.

Within-delay zero-order correlations were also non-significant (6h: r = -0.12, p = 0.49; 24h: r = -0.12, p = 0.44; 48h: r = +0.15, p = 0.32), as were the corresponding partial correlations controlling for memory strength (6h: r = -0.13; 24h: r = -0.14; 48h: r = +0.20, p = 0.19). At 48 hours the numerical sign was positive, but the magnitude was small and the effect did not approach significance.

We ran a parallel analysis for accuracy in the perceptual discrimination task (all trials), which does not require the minimum-trial exclusion used for the bias index and so draws on the full sample (Fig. 6b; n = 177; 6h n = 56, 24h n = 64, 48h n = 57). In a multiple regression predicting perceptual accuracy from cue generalisation while controlling for delay group and semantic recall accuracy, the coupling coefficient was negative but did not reach conventional significance (β = -0.124, SE = 0.067, t(172) = -1.86, p = .065, 95% CI [-0.256, +0.008]). The estimate was similar but further attenuated when memory strength was excluded from the model (β = -0.100, p = .238), and the cue-generalisation × delay interaction did not significantly improve model fit, F(2, 170) = 2.59, p = .078.

Within-delay correlations showed that this marginal overall association was driven almost entirely by the 24-hour delay: cue generalisation and perceptual accuracy were significantly negatively correlated at 24h (r = -.297, p = .017; partial r controlling for memory strength = -.340, p = .006), but not at 6h (r = +.084, p = .540; partial r = -.005, p = .970) or 48h (r = -.035, p = .796; partial r = -.029, p = .831).

Taken together with the null result for the Typicality Bias Index, these findings suggest that perceptual accuracy and cue generalisation were not coupled across the sample as a whole, with a significant negative relationship emerging only at the intermediate 24-hour delay where participants who generalised more from the more-typical cue also tended to show somewhat lower perceptual accuracy at that time point specifically. Because this relationship was absent at both the earliest (6h) and longest (48h) delays and the overall interaction test was only marginal, we treat it as a locally emerging pattern rather than strong evidence of a delay-general coupling between the two measures.

## Discussion

Our results reveal a feature of adaptive memory. As episodic memories consolidate, retrieval increasingly exploits category structure, while the concrete perceptual content of the original episode remains accessible. The benefits of cue generalisation occur without the cost of losing concrete details. Over time, the originally encoded exemplar, which is the most effective trigger shortly after learning, comes to share its privileged status with never-encoded but more typical exemplars. And when perceptual discrimination failed, reconstruction was consistently biased towards category-typical alternatives, a pull already evident at the shortest delay. If these changes were the expression of a single, unified representation sliding from an episodic towards a schematic state, they should constitute a coherent package: cue generalisation, typicality-biased reconstruction, and the erosion of concrete detail should co-occur. They did not. We found robust cue generalisation and a reliable bias towards category-typical reconstructions, but no evidence that perceptual detail eroded across delay, and neither the typicality bias nor perceptual accuracy reliably covaried with cue generalisation across participants. Consolidation therefore appears to act not along a single episodic-to-schematic dimension, but on partially separable components of the same memory trace.

Tulving and Thomson’s encoding-specificity principle holds that retrieval succeeds to the extent that the information available at test overlaps with what was encoded (Tulving & Thomson, 1973), and later work on cue-dependent retrieval and context reinstatement reinforced the view that effective cues are those that are both available and diagnostic of the target (Nairne, 2002; Smith & Vela, 2001; Watkins & Watkins, 1975). The principle is sometimes read statically, as though the effective cues were fixed by what was present at study. Yet cue effectiveness has been shown to depend on a cue’s diagnosticity relative to competitors (Nairne, 2002) and suggested to depend on the current state of the memory representation itself (Linde-Domingo & Kerrén, 2025). Our findings support this dynamic reading as an extension of encoding specificity rather than a contradiction of it. The most effective cue changes across delay because the trace it must contact changes across delay. At 48 hours, a never-encoded but more typical exemplar supported retrieval as well as the original studied image. Less-typical exemplars, despite sharing the same verbal label, remained numerically less effective and did not show the same clear evidence of convergence. This dissociation is difficult to reconcile with an unchanging trace accessed through a fixed optimal cue, but follows naturally if consolidation reweights the categorical components of the representation, so that access increasingly proceeds through schema-like structure (Gilboa & Marlatte, 2017). Notably, our preregistered prediction anticipated that more-typical cues would eventually outperform the originally encoded cue. Instead, we observed convergence. This pattern may itself be theoretically informative. If memory transformation broadens access through category-level information while episode-specific information remains available, there is little reason for an adaptive memory system to lose the effectiveness of the original retrieval route. Generalisation may therefore expand, rather than replace, the routes through which an episode can be accessed.

When we examined whether the visual details of the associated item remained accessible over time, discrimination of the exact encoded exemplar remained robustly above chance and its accuracy was largely stable across delay, even as response times slowed; with the one exception being a modest decline in the rejection of novel-concept lures by 48 h. Durable recognition of object detail over long retention intervals has been reported before (Brady et al., 2008; Konkle et al., 2010), so above-chance discrimination at 48 h is not in itself surprising. What is theoretically informative is the company this preserved detail keeps. Although concrete exemplar recognition might be thought to be solvable through image familiarity alone, two features of our data argue otherwise. A purely familiarity-based strategy predicts no influence of the cue; yet response speed was systematically graded by which exemplar had served as the retrieval cue, revealing an associative component to the decision. Converging with this, trial-level accuracy in the perceptual discrimination task was predicted by memory strength estimated from an earlier semantic recall test of the same association. Such dependency cannot be easily explained by a familiarity signal, since recognising the target image should not depend on how well its associate had been retrieved from a different cue. Together, these features indicate that exact-exemplar selection engaged the stored association rather than image familiarity alone.

A complementary picture emerges from the errors themselves in the perceptual discrimination task. When participants failed to select the exact target, their mistakes were not evenly distributed across alternatives but, instead, were asymmetrically drawn towards the more typical exemplar of the target’s category. This asymmetry was already robust at the shortest delay, when memory was strongest, and persisted across delays, suggesting that such bias shapes memory reconstruction from the start. Such directional distortion is precisely what reconstructive accounts of memory predict. Category-adjustment (Huttenlocher et al., 2000), Bayesian (Hemmer & Steyvers, 2009a; Spens & Burgess, 2024), and fuzzy-trace (Brainerd & Reyna, 2002) models converge in predicting that when item-specific information is insufficient, memory judgements become increasingly influenced by generalised or category-level information, producing systematic shifts towards more prototypical content. It is important to mention that such typicality bias also appears in the target-absent trials, on which all three alternatives were presented from a concept that had never been studied and the correct response was to reject them all. Here too, errors favoured the more typical exemplar, and this preference grew with delay. Because no episodic representation of these images existed, the bias on these trials cannot reflect the transformation of a stored trace. What possibly remains is a decision-level tendency to endorse typical items when episodic evidence is insufficient based on plausibly greater fluency and familiarity of such items. But also, that errors on target-absent trials are still pulled towards typical exemplars fits a broader view of reconstructive retrieval as integration of stored evidence with plausible, structure-consistent content, a process that can yield systematic distortions or even false memories when the veridical trace is unavailable (Loftus, 2005; Schacter et al., 2011). Two components thus appear to combine in our error data: a general, uncertainty-scaled reliance on category structure, which grows with delay, and a trace-specific prototype attraction, which is present at the earliest delay and changes comparatively little thereafter.

This category-structure bias raises a question: does this drift towards typical items simply happen when memories are weak? Our data do not support this reduction. Trial-based memory strength (indexed by accuracy, confidence and speed of semantic recall) clearly predicted perceptual accuracy. However, it was unrelated to whether an error, once made, drifted towards the more or less typical examples. Memory strength and reconstructive bias therefore behaved as separable dimensions: one determined access to the veridical trace and the other one the bias of reconstruction when retrieval fails.

Such separation aligns well with a growing body of work supporting the idea that episodic memory is rather a multidimensional content whose features can vary independently (see Heinen et al., 2025; Horner, 2026). For instance, Berens and colleagues (2020) and more recently Raz Groman and Sadeh (2025), found that forgetting reduces the accessibility of a memory without necessarily affecting its precision, dissociating the retrieval from the fidelity of what is retrieved. Our results add a third, orthogonal property to this picture: the direction of reconstructive bias. Here we show that such components are not tightly coupled to availability. An item can be hard to retrieve without its errors being any more typical-biased, and such a bias can grow with delay without any accompanying change in how strongly items are accessed. This does not contradict accounts in which imprecision invites schema-based filling-in (Hemmer & Steyvers, 2009a; Huttenlocher et al., 2000; Ramey, 2026; Schacter et al., 2011); rather, it refines them, by indicating that the degree to which the prior is weighted is not fixed by episodic strength alone but varies partly independently, here as a function of elapsed time.

A similar conclusion emerges from a different angle. If episodic memory were a unified phenomenon, expanding the effectiveness of retrieval cues to never-encoded information should be linked to a potential specificity loss. If that were the case, participants who showed a stronger cue generalisation towards never-encoded but more typical cues should also have a higher typicality bias in errors. However, cue generalisation did not contribute to typicality bias or perceptual accuracy. We are cautious in interpreting a null, and the present sample bounds rather than eliminates the possibility of a modest association. But taken together with the trial-level dissociation, the pattern is difficult to reconcile with a strictly one-dimensional account. It fits more naturally with the view that consolidation acts on partially separable components of a memory.

We opened with a memory that had to survive being altered: a portrait recognised despite the changes made to it, requiring both tolerance of a transformed cue and retention of the details that identified the original. The present results suggest that episodic memory resolves this demand without trading one capacity for the other, showing its adaptive nature. Over hours and days, the originally encoded exemplar lost its privileged status as a retrieval probe to never-encoded but more typical alternatives, while the perceptual content of the associated episode remained discriminable, and reconstruction, when retrieval failed, was drawn towards the category-typical information. Critically, these changes move rather independently. Memory strength predicted whether the target was recovered but not the direction of errors, and the generalisation of effective cues was not reliably coupled, across individuals, to either the preservation of detail or the typicality bias. Memory transformation therefore appears to act not as a single change from the episodic towards the semantic, but on partially separable components of the same trace: it extends the routes by which an episode can be reached while leaving its specific content accessible to those who reach it.

### Limitations

Some features of the design limit interpretation. Retrieval tasks were presented in a fixed order so that the perceptual discrimination task, which necessarily re-exposes participants to the encoded image, could not contaminate the measure preceding it. Also, the categorisation task, answered first on every trial, yielded an estimate of retrieval speed not affected by previous retrieval attempts. Although such fixed order has some benefits, assigning a single retrieval measure to each trial in future studies could provide a cleaner estimate, although at a substantial cost in power.

Delay was manipulated between participants. Although a within-participant design would allow greater sensitivity, repeated testing of the same associates would alter the memories under study (Antony et al., 2017; Ferreira et al., 2019; Hupbach et al., 2007; Lifanov et al., 2021; Nader & Hardt, 2009; Roediger & Butler, 2011). Alternatively, testing different associations for each participant at each delay would be an option for future studies but, also, at a cost in power. The current between-participant design avoids this, and encoding-phase performance did not differ across delay groups. However, it means that delay effects are group comparisons, and that recruitment proceeded sequentially by delay condition.

Our delay conditions also differ in more than elapsed time. The 24-h and 48-h groups are expected to sleep between encoding and test, the 6-h group in many cases did not. So the groups differ additionally in waking interference, circadian phase, and time of day at test. Recent work indicates that sleep can preserve item-level representations while strengthening category-level structure (Schapiro et al., 2017), a combination resembling the pattern reported here and it is expected to have a key role in memory consolidation (Diekelmann & Born, 2010; Rasch & Born, 2013). Although recent evidence suggests that memory generalisation can also increase across delay independently of post-encoding sleep (Lu et al., 2026; Lutz et al., 2026). The present design does not distinguish between effects derived from sleep from elapsed time so we therefore treat it as one candidate mechanism rather than an explanation. A within-subject retest design, nap-versus-wake manipulation, or polysomnography follow-up could address this directly.

The present finding raises questions that the data cannot answer. Convergence between encoded and more-typical cues emerged at the delay where categorisation performance was lowest, raising the possibility that convergence could reflect compression towards a common floor. The differential decline rate, the selectivity of generalisation for the more-typical but not the less-typical cue, and above-chance performance throughout argue against a pure floor account, but do not fully exclude it. Manipulating encoding strength independently of delay would help to establish whether the effect we report is a general property of consolidating memories or a feature of a particular region of the performance range (see for instance how poorly encoded information overgeneralised in the rodent literature; Zinn et al., 2020).

A related boundary condition concerns cue diagnosticity (Nairne, 2002). Our design preserved a one-to-one relationship between each cue concept and a single associated target. Thus, even after generalisation, a category-matching cue remained diagnostic of one association. If several exemplars from the same concept were associated with different targets, broader cue efficacy could instead reduce diagnosticity and increase retrieval competition or cue overload (Watkins & Watkins, 1975). Future work should therefore test whether the generalisation of cue efficacy persists under these conditions.

A second set of questions concerns the neural patterns associated with these results. On one side, the behavioural dissociation invites investigation of the neural mechanisms behind cue generalisation: if cue-generalisation occurs while visual details are preserved, it is possible that the two carry separable neural signatures. Although these limitations and future questions remain, these findings offer valuable results for understanding the multidimensional and adaptive nature of episodic memory.

## Data availability

The data that support the findings of this study are available at https://gin.g-node.org/m.delmarco/prototypical-object-paradigm

## Code availability

The code to reproduce the results of this study is available at https://gin.g-node.org/m.delmarco/prototypical-object-paradigm

## Acknowledgments

C.G.G. was supported by Project PID2023-149428NB-I00 funded by MCIN/AEI/10.13039/501100011033 and by FEDER, EU, and Grant RYC2021-033536-I funded by MCIN/AEI/10.13039/501100011033 and by the European Union NextGeneration EU/PRTR. J.L.D. was supported by Project PID2023-151104NA-I00 funded by MCIN/AEI/10.13039/501100011033 and by FEDER, EU, and Grant RYC2021-033940-I funded by MCIN/AEI/10.13039/501100011033 and by the European Union NextGeneration EU/PRTR. The Mind, Brain and Behavior Research Center receives funding from grants CEX2023-001312-M by MICIU/AEI/10.13039/501100011033 and UCE-PP2023-11 by the University of Granada. The funders had no role in the study design, data collection and analysis, decision to publish or preparation of the manuscript. This manuscript is part of the PhD thesis of M.D.

## Author contributions

J.L.D., M.D., and C.G.G. contributed to the conceptualization and methodology of the study, and to writing, review, and editing of the manuscript. J.L.D. was responsible for project administration, funding acquisition, formal analysis, visualization, and writing of the original draft. M.D. contributed to data curation, formal analysis, investigation, visualization, and writing, review, and editing. C.G.G. contributed to conceptualization, methodology, and writing, review, and editing.

## Competing interests

The authors declare no competing interests.

## Notes

https://gin.g-node.org/m.delmarco/prototypical-object-paradigm

### Competing Interest Statement

The authors have declared no competing interest.

