## Supplementary Figures for "Episodic memory adaptively expands retrieval through typicality-guided transformation while concrete details remain accessible"

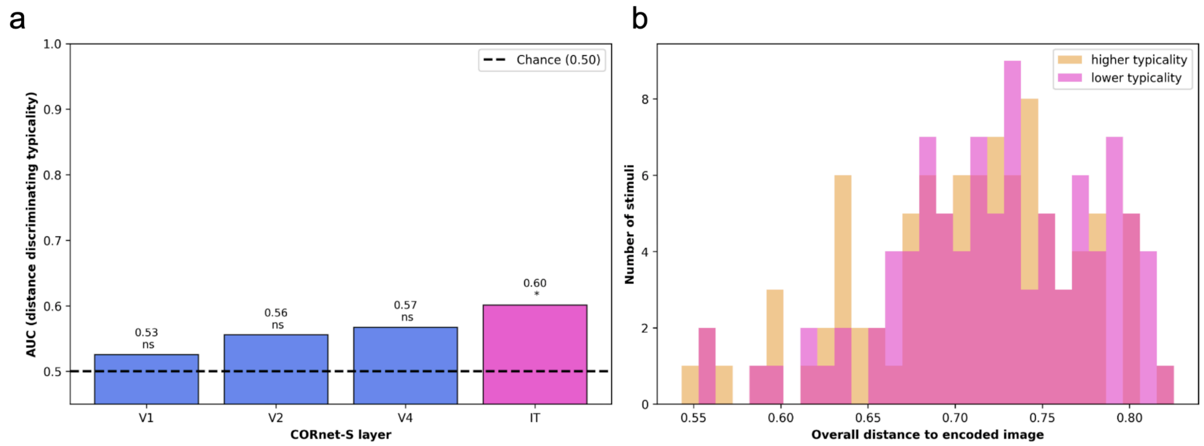


**Figure S1. Decodability of cue typicality from visual distance to the encoded image.** We tested whether the more versus less-typicality cue manipulation could be explained by systematic differences in visual similarity to the encoded exemplar. Image representations were extracted from the V1, V2, V4, and IT layers of the CORnet-S deep neural network (Kubilius et al., 2019), and representational similarity analysis was used to calculate the distance between each encoded image and its corresponding higher and lower-typicality cues. Because both typicality and DNN distance were fixed stimulus properties, analyses were performed at the stimulus level. The analysis included 94 object concepts with complete DNN distances, each contributing one higher and one lower-typicality observation, for a total of 188 stimulus-level observations. Eighteen of the original 112 concepts were excluded because DNN distance estimates were unavailable. **a.** Separability between higher and lower-typicality cues was tested independently for each CORnet-S layer. The plotted AUC was derived from the Mann–Whitney statistic and can be interpreted as the probability that a randomly selected lower-typicality cue was farther from its encoded image than a randomly selected higher-typicality cue. An AUC of .5 therefore indicates no systematic association between cue typicality and distance to the encoded image. As a complementary decoding analysis, a single-feature logistic-regression classifier was fitted separately for each layer and evaluated using five-fold stratified cross-validation. Statistical significance was assessed by comparing the cross-validated ROC-AUC against a null distribution generated from 1,000 label permutations; asterisks above the bars indicate the permutation-test results. Distance did not reliably discriminate cue typicality in V1, V2, or V4 (AUCs = .53, .56, and .57; permutation *ps* = .404, .156, and .079, respectively). Typicality was decodable from IT-layer distance (AUC = .60, permutation $p$ = .019), indicating some association between typicality and higher-level CORnet-S information, but little evidence that the manipulation was driven by low-level visual similarity. **b.** Overlapping distributions show the overall distance to the encoded image for higher and lower-typicality cues, calculated by averaging distances across the four CORnet-S layers. The substantial overlap illustrates that the two cue types were broadly similar in their visual proximity to the encoded exemplar, despite the modest separation observed at the IT layer.

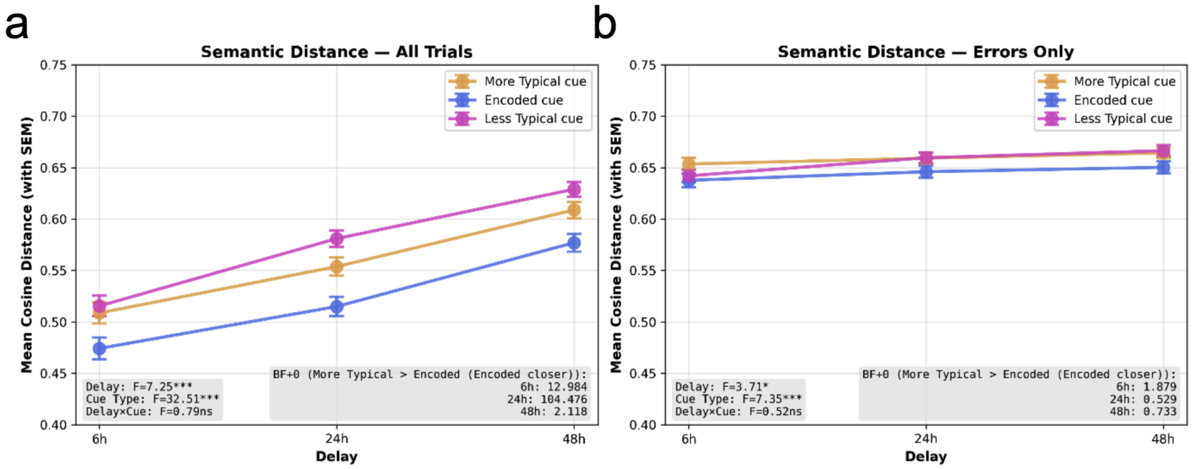


**Figure S2. Semantic distance in the semantic recall task.** Mean cosine distance between participants' typed responses and the correct target label, shown as a function of delay and type of cue. Lower values indicate responses that were semantically closer to the target. **a**, Semantic distance across all trials. Responses became less semantically close to the target with increasing delay, $F(2, 174) = 7.25$, $p < .001$, $\eta_p^2 = .077$, and differed reliably by cue type, $F(2, 348) = 32.51$, $p < .001$, $\eta_p^2 = .157$. The delay × cue interaction was not significant, $F(4, 348) = 0.79$, $p = .529$, $\eta_p^2 = .009$. Across delays, responses were closest to the target following encoded cues, followed by more-typical and less-typical cues. Bayesian comparisons between encoded and more-typical cues supported an encoded-cue advantage at 6 h and 24 h, and were weaker at 48 h ($BF_{+0}$ = 12.98, 104.48, and 2.12, respectively). **b**, The same analysis restricted to incorrect trials only. Semantic distance still increased with delay, $F(2, 173) = 3.71$, $p = .026$, $\eta_p^2 = .041$, and differed modestly by cue type, $F(2, 346) = 7.35$, $p < .001$, $\eta_p^2 = .041$, with no delay × cue interaction, $F(4, 346) = 0.52$, $p = .718$, $\eta_p^2 = .006$. However, cue-related differences were much smaller when only errors were considered, and Bayesian comparisons did not provide clear evidence for an encoded-cue advantage over more-typical cues ($BF_{+0}$ = 1.88, 0.53, and 0.73 at 6 h, 24 h, and 48 h, respectively). Thus, the stronger cue-type effect observed across all trials appears to be driven mainly by differences in exact or near-exact target retrieval, rather than by large differences in the semantic proximity of incorrect responses. Error bars indicate ±1 SEM.

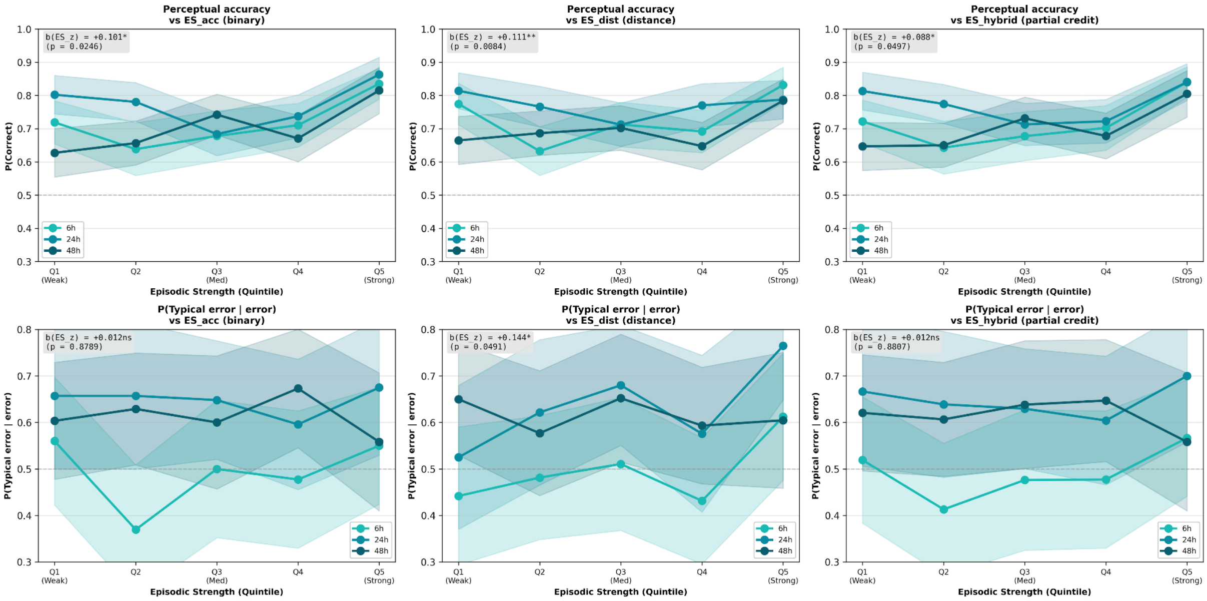


**Figure S3. Robustness of episodic-strength effects across alternative operationalisations.** Perceptual accuracy and typicality bias were re-analysed using three alternative estimates of item-level episodic strength derived from the semantic-cued recall task. The accuracy-based index, as reported in the main text, treated semantic recall as binary and combined semantic accuracy with confidence and response speed: ES_acc = semantic accuracy + 0.5 × z(confidence) − 0.5 × z(log RT). The distance-based index replaced binary accuracy with graded semantic precision, using the negative z-scored semantic distance between the typed response and the target label: ES_dist = −z(semantic distance) + 0.5 × z(confidence) − 0.5 × z(log RT). The hybrid index combined both approaches by assigning full credit to correct semantic responses and partial credit to incorrect responses according to their semantic proximity to the target: ES_hybrid = graded semantic accuracy + 0.5 × z(confidence) − 0.5 × z(log RT), where incorrect responses received higher partial credit when they were semantically closer to the target. All analyses used the same complete-case trial samples across the three indices within each dependent variable, an approach that reduced the number of trials for the ES_acc here compared to the results shared in the main text. The top row shows perceptual accuracy as a function of episodic-strength quintile, separately for each delay group. Across all three operationalisations, stronger episodic strength predicted higher perceptual accuracy in logistic GEE models: ES_acc, $b$ = 0.101, $p$ = .025; ES_dist, $b$ = 0.111, $p$ = .008; ES_hybrid, $b$ = 0.088, $p$ = .050. The bottom row shows the probability that a perceptual error was directed toward the more-typical exemplar. This relationship was absent for the binary and hybrid indices, and only weakly present for the distance-based index: ES_acc, $b$ = 0.012, $p$ = .879; ES_dist, $b$ = 0.144, $p$ = .049; ES_hybrid, $b$ = 0.012, $p$ = .881. Thus, the central conclusion was robust to how episodic strength was estimated, where stronger item-level memory predicted whether participants could identify the target, whereas the direction of perceptual errors was not consistently explained by episodic strength. Shaded bands denote 95% confidence intervals; dashed horizontal lines mark 0.5, corresponding to no directional typicality bias in the lower panels.
